# A T-cell intrinsic Role for *APOL1* Risk Alleles in Allograft Rejection

**DOI:** 10.1101/2025.07.14.664793

**Authors:** John Pell, EM Tanvir, Zeguo Sun, Irene Chernova, Anand Reghuvaran, Soichiro Nagata, Mateus Guerra, John Choi, Soltan Al Chaar, Hiroki Mizuno, Ke Dong, Xin Tian, Reika Ishibe, Barbara Franchin, Paolo Cravedi, Ashwani Kumar, Gabriel Barsotti, Hongmei Shi, Bony De Kumar, Wenzhi Song, Cijiang He, Jordan S Pober, Stefan Somlo, Ian W Gibson, Waldemar Popik, Zhongyang Zhang, Joseph Craft, Jamil Azzi, Naoka Murakami, Shuta Ishibe, Peter Heeger, Madhav C Menon

## Abstract

African Americans have an increased risk of kidney disease due to exonic variants in Apolipoprotein-L1 (G1 and G2). These prevalent variants have also been linked with kidney rejection, but outside of association with African ancestry, underpinning causal mechanisms are unknown. We investigated T-cell function using transgenic mice with physiologic expression of wild type (G0-), G1-, or G2-APOL1. Mice with variant APOL1 showed greater CD8+T-cell activation with expansion of a central memory (TCM) subset. Stimulated G1-CD8+T-cells showed enhanced proliferation and cytokine production, which reversed with APOL1 inhibition. In MHC-mismatched cardiac transplants, G1-mice demonstrated greater CD8+T-cell infiltration and reduced survival. Bulk transcriptome of G1-CD8+T-cells, and single-cell transcriptome of graft infiltrating TCMs, showed enrichment of canonical T-cell receptor (TCR) pathways including Ca^2+^-signaling. G1-CD8+T-cells demonstrated baseline ER-Ca^2+^ depletion followed by sustained increases in cytosolic-Ca^2+^ upon TCR stimulation. G1-CD8+T-cells were more sensitive to Ca^2+^ chelation, or store-operated Ca^2+^ entry inhibition, and relatively resistant to calcineurin antagonism vs. G0-CD8+T-cells. Analogously, in a kidney transplant cohort, APOL1-variant recipients developed rejection when they had elevated peripheral TCMs before transplantation and despite significantly higher tacrolimus levels vs G0/G0-AAs with rejection. In summary, we unravel an excitatory T-cell intrinsic mechanism for APOL1 exonic variants, causally linking them with kidney rejection.

## INTRODUCTION

African Americans (AAs) are at disproportionate risk of progressive kidney disease and focal segmental glomerulosclerosis (FSGS) due to the carriage of exonic variants (G1 and G2) in the Apolipoprotein L1 gene (*APOL1)* (1, 2). G1-, G2-APOL1 are seen exclusively within AAs or patients with admixed African ancestry, whereas all other ancestries carry the major allele (G0). APOL1-variants emerged as part of an “evolutionary arms race” against *Trypanosoma brucei spp* which causes African trypanosomiasis [reviewed in (3)]. Following the seminal discovery of FSGS-associated APOL1-variants (APOL1-RV used hereon for 1- or 2-risk variants) (4), outstanding contributions have improved understanding of APOL1-FSGS. APOL1 is present only in some primates and apes, limiting *in vivo* models (5). Elegant murine data showed that podocyte-specific, over- expression of APOL1-RVs, induced FSGS compared to G0 (6). The mechanism of APOL1-induced cytotoxicity in podocytes is less certain despite detailed studies using in vitro (7), non-mammalian (8) or rodent models (9-11). APOL1 is a monovalent cation channel localized to organelle-(8) and plasma-membranes (12, 13), and increased channel activity in APOL1-RVs (vs G0-APOL1) is likely a key mechanism for cytotoxicity in kidney epithelial cells (11, 14), which is targeted by pharmacotherapeutics (15). Clinical data from kidney transplantation showed that the presence of APOL1-RVs *in kidney donors* increases the risk of death-censored allograft loss (DCAL) (16), also supporting a kidney epithelial cell-centric role of APOL1-RVs in disease risk.

Provocative data from our laboratory suggested that the presence of APOL1-RVs *in kidney recipients* are associated with increased T-cell mediated rejection (TCMR) and DCAL (17), independent of donor-APOL1. Similar findings associating APOL1-RVs in kidney recipients with TCMR and DCAL have been reported by two independent studies (18, 19). Interestingly, recipient APOL1-RVs associated with graft outcomes in an additive manner, i.e., every copy associated with increased risk (17, 19). Together, these data raise the prospect of a hitherto unexplored role for APOL1-RVs in T-cells which are central to TCMR. However, even well-conducted epidemiologic studies cannot fully uncouple causation from association, especially since AAs (who carry APOL1-RVs) are also at higher risk of rejection from multiple factors (19-21). Therefore, defining the role of APOL1-variants in T-cells using mechanistic models has considerable importance for kidney transplants due to the high prevalence of this variant among AAs (∼40% AAs have at least one copy of risk-variants).

To investigate the role and mechanism of APOL1-RVs in T-cells, we generated bacterial artificial chromosome transgenic (BAC-Tg) mice with the human *APOL1* promoter for G0, G1, and G2. BAC-Tg models result in physiologic levels of *APOL1* as BAC inserts include *cis*-regulatory sequences to mirror endogenous transcription. Using this model, we report that APOL1-RV BAC-Tg mice show T-cell activation (17), and expansion of a T-central memory (TCM) subset. We then performed a series of mechanistic experiments incorporating inter-related translational human studies to identify a role and unravel the mechanism of APOL1-RVs in the context of T-cell activation, allograft rejection and graft survival.

## RESULTS

### Generation and phenotyping of APOL1-BAC-Tg animals

APOL1-BAC transgenic mice (G0, G1, G2-BAC-Tg) were generated as detailed (Figure 1A; see methods). Mice were obtained in a hybrid genetic background (G0-, G1-, G2-hybrid;), and only mice with ∼2 copy numbers of APOL1 BAC transgenes were used for experiments (Supplemental Figure 1A). The expression of *APOL1* mRNA, and APOL1 protein were confirmed in splenocyte lysates (Supplemental Figure 1, B and C).

**Figure 1.**
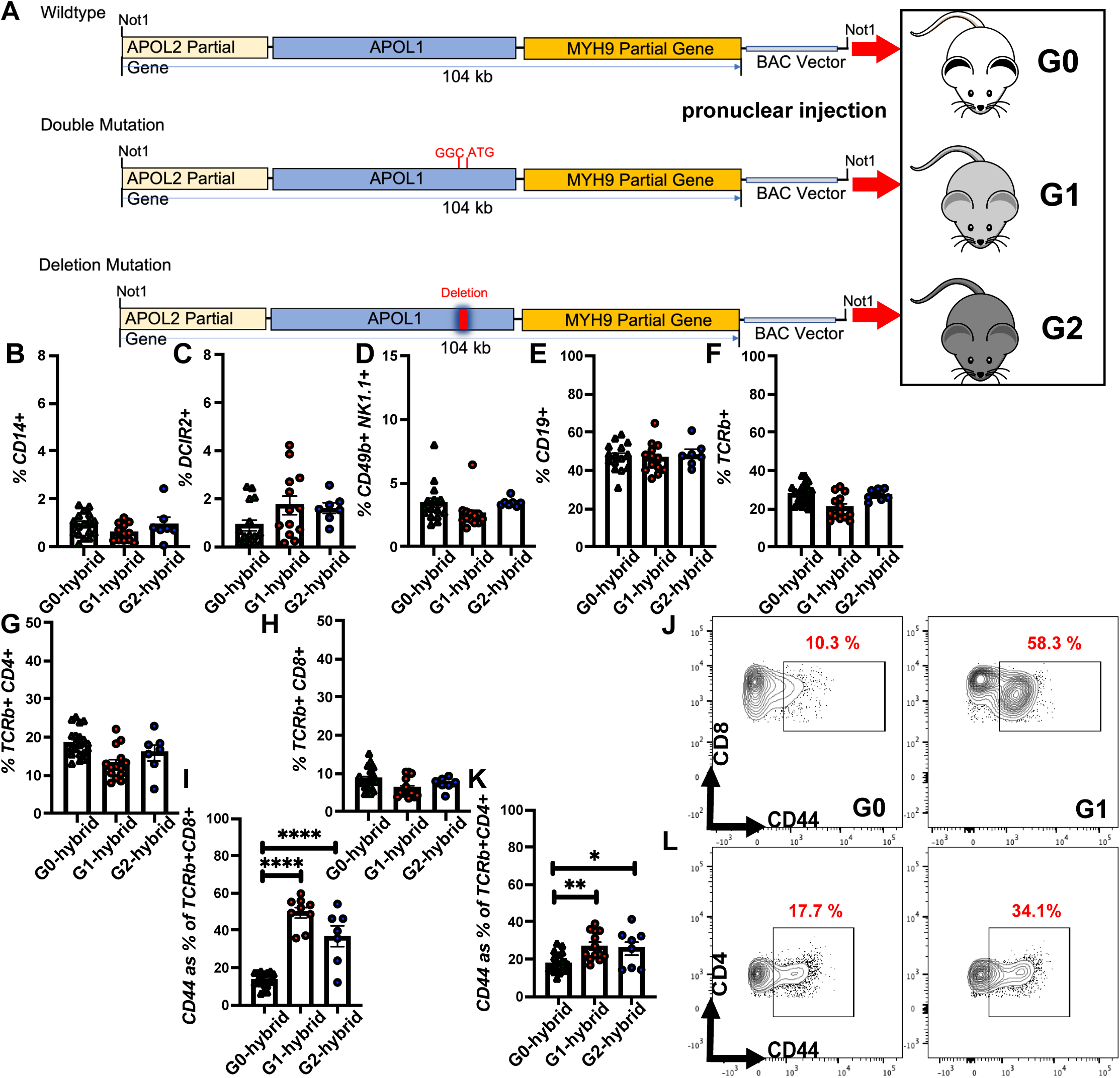
APOL1-BAC transgenic mice development and baseline T-cell phenotypes. (A) APOL1-BAC-Tg mice were developed by pronuclear injection using a 104 kb BAC into oocytes to generate G0-, G1-, and G2-BAC-Tg lines. (B-H) Baseline FACS analysis of splenocytes compares percentages of innate immune cells (CD14+, DCIR+, CD49b+NK1.1+), as well as adaptive immune cells (B-cells (CD19+), T-cells (TCRb+), CD4+ and CD8+T-cells) between the three lines. (I-L) Bar graphs of percentages of (I) CD8+CD44+ and (K) CD4+CD44+ cells among T-cells in spleens of BAC-Tg lines at baseline. (J) and (L) are representative flow plots of data in (I) and (K), respectively. The bar graphs display the mean ± SEM of n ≥ 7 mice. *p<0.05, **p<0.01, ****p <0.0001. Individual mice are represented by circles (G1, G2) and triangles (G0).

We first studied splenocyte fractions by flow cytometric analysis (FACS) in G0-, G1-, G2-hybrid mice (of both sexes, 8-10 weeks old) to understand baseline immunologic phenotypes [n ≥ 7 mice each]. Total splenocyte numbers were not significantly different between BAC-Tg lines (Supplemental Figure 1D). The proportions of major innate immune- (dendritic cells (DCIR+), monocyte-macrophages (CD14+), NK cells (NK1.1+)) and adaptive-immune cells- (B-cells (CD19+) and T-cells (TCRb+), CD4+ and CD8+T-cells) were similar across the genotypes (Figure 1, B-H and Supplemental Figure 1E for gating strategy). Based on our prior human data showing T-cell activation in variant-APOL1 individuals (17), we focused on T-cell phenotype in BAC-Tg splenocytes using a characterized flow cytometric panel for T-cell subsets (22). Interestingly, CD8+ and CD4+ T-cells showed a significantly increased proportion of activation marker CD44 in G1-, G2-hybrid mice vs. G0-hybrid at baseline (Figure 1, I-L), with greater increases among CD8+ CD44+T-cells vs. G0 than among CD4+T-cells (Figure 1, I vs K). CD8+CD69+ and CD4+CD69+T-cell proportions (early activation or tissue-resident marker), were not significantly different between G0-, G1- or G2-hybrid mice at baseline (Supplemental Figure 1, F-I). In homeostasis at 12 weeks, APOL1-RV BAC-Tg mice showed no evidence of azotemia or albuminuria vs G0-BAC-Tg (Supplemental Figure 1, J and K). APOL1-RV BAC-Tg mice aged up to 1 year also showed no weight loss or alopecia (not shown).

### Variant APOL1 mouse splenocytes demonstrate increased CD8+ and CD4+T-cell activation at baseline in adult animals

Since T-cell phenotype is influenced by genetic background (23), we backcrossed our hybrid BAC-Tg lines into a C57Bl/6J background to generate G1- and G0-BAC-Tg B6 lines (called G0 and G1 hereon) (Supplemental Figure 2A). APOL1-copy numbers were maintained through six backcrosses and showed no dilution of gene dose (Supplemental Figure 2B). We used G2-hybrid-Tg mice to validate selected findings. Among CD8+ T-cells, CD44+ proportions remained higher in backcrossed G1 mice vs G0 (Figure 2A). CD8+CD44+T-cell proportions were significantly higher in G1- than WT-B6 splenocytes, whereas G0-CD8+CD44+T-cell proportions were similar to WT-B6 (Figure 2A), suggesting gain-of-function with G1. Among CD8+CD44+T-cells, significantly higher percentages of CD8+CD44+CCR7+T-cells (TCMs) were observed (Figure 2, B and C), while CD8+CD44+CCR7-T-cells were not increased (Figure 2D). Expansion of the TCM subset in G1-BAC-Tg mice was not identifiable at weaning (4 weeks old) but was observed in all older mice (Supplemental Figure 2C) and in both sexes. CD8+CD44+CXCR3+T-cells were increased in G1-BAC-Tg (Figure 2, E and F). Interestingly, CD8+CD44+KLRG1+T-cells were significantly reduced in G1 mice vs. G0 (Figure 2, G and H). Intracellular transcription factor staining showed significant increases in memory T-cell marker Eomes+ (Figure 2, I and J) and stemness marker Tcf1+ (Figure 2, K and L) in G1-CD8+CD44+T-cells. The proportions of Blimp1+, Tbet+, and Rorgt+ were not different between G0- and G1-CD8+CD44+T-cells (Supplemental Figure 2, D-F). G1-CD8+ T-cells also showed increased levels of cytotoxicity marker Perforin vs. G0 (Figure 2, M and N).

**Figure 2.**
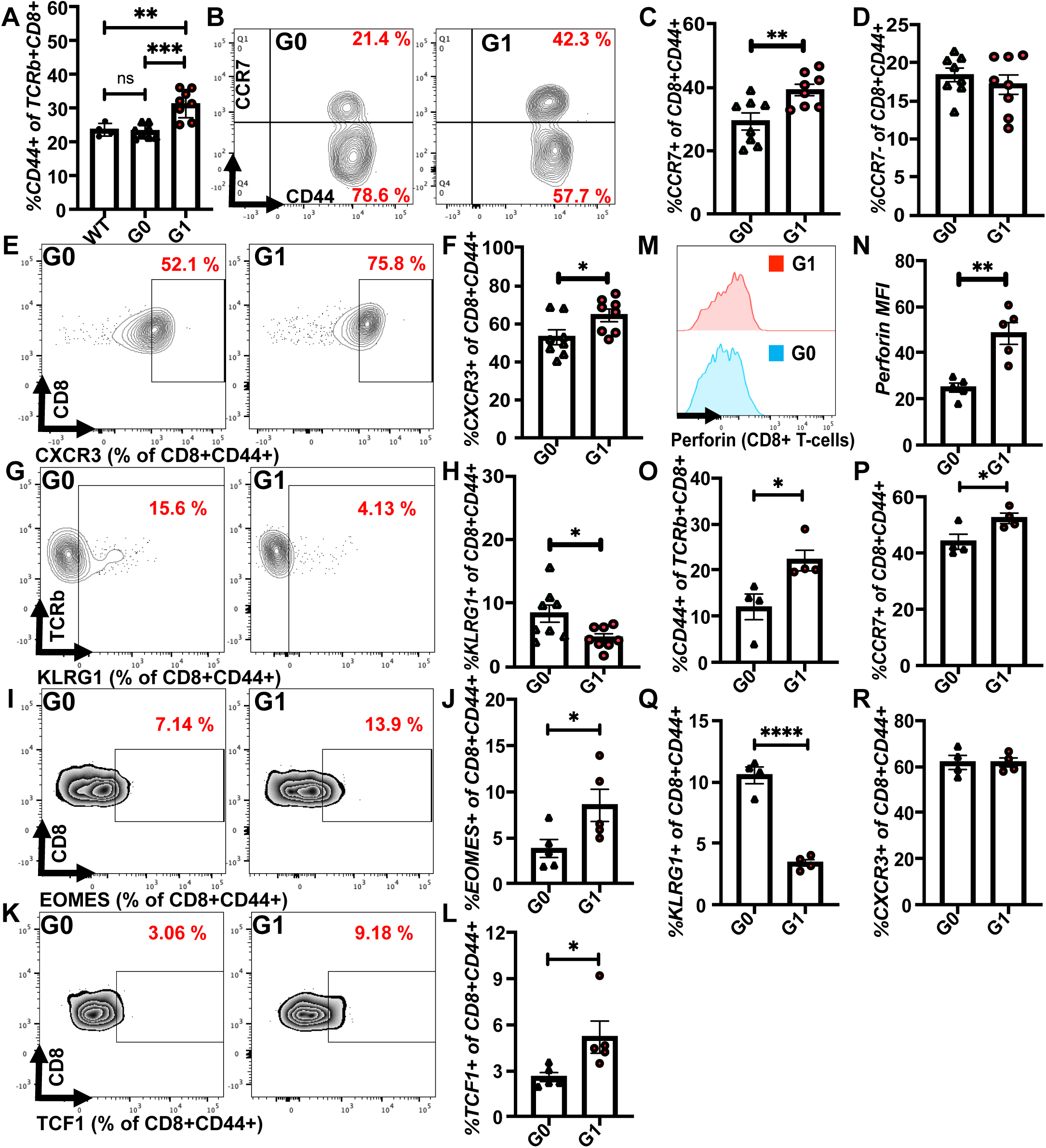
Baseline activation and polarization of CD8+ and CD4+ T-cells in backcrossed variant APOL1-BAC-Tg mice. (A) Proportions of CD44+CD8+T-cells in G1, G0 and WT-C57Bl/6J mice. (B) Representative flow plots of CD8+CD44+T-cells gated for CCR7+ cells to identify CD8+CD44+CCR7+T-cells (T-central memory or TCMs). Quantified proportions of (C) TCMs and (D) CD8+CD44+CCR7-T-cells [G1 vs. G0; n>5 mice each]. (E-H) Representative flow plots and quantifications of the proportion of CD8+CD44+T-cells gated for (E and F) CXCR3+, (G and H) KLRG1+cells between G0- and G1-mice (n>5 mice each). (I-L) Representative flow plots and quantifications of cell proportion from intracellular staining of CD8+CD44+T-cells gated for the transcription factors (I and J) EOMES and (K and L) TCF1 between G0- and G1-mice (n=5 mice each). (M) and (N) representative flow plot and quantified mean fluorescence intensity of perforin, respectively, in G0 and G1 mice. Mesenteric lymph nodal T-cells were evaluated for proportions of (O) CD8+CD44+T-cells, (P) TCMs (CD8+CD44+CCR7+T-cells) (Q) CD8+CD44+KLRG1+T-cells and (R) CD8+CD44+CXCR3+T-cells (n=4 mice each). The bar graphs display the mean ± SEM of n≥7 mice. *p<0.05, **p<0.01, ***p<0.001, ****p<0.0001. Circles/triangles represent individual mice.

Among CD4+T-cells, significantly increased CD4+CD44+T-cells were also seen in G1-BAC-Tg (Supplemental Figure 2G); however, CXCR3+ or CD4+CD44+CCR7+T-cell expansion was not observed (Supplemental Figure 2, H and I). G1-CD4+CD44+T-cells also showed significantly higher Tcf1 vs G0 (Supplemental Figure 2J). Eomes+, Blimp1+, Tbet+, and RORgT+ cells were not different between G0- and G1-CD4+CD44+T-cells (Supplemental Figure 2, K-N). Th2 fractions (CD4+GATA3+) were also not different between G0 and G1 lines (Supplemental Figure 2O). Tregs (CD4+FoxP3+T-cells) were significantly reduced in pre-backcrossed G1-spleens (vs G0) but were similar to G0-BAC-Tg after backcrossing (Supplemental Figure 2, P and Q).

Analysis of mesenteric lymph nodes also confirmed increased CD8+CD44+T-cells, TCM expansion and decreased CD8+CD44+KLRG1+T-cells, but showed similar CD8+CD44+ CXCR3+T-cells, in G1 vs G0 (Figure 2, O-R). No significant differences in activation or subset expansion were seen among mesenteric lymph nodal CD4+T-cells in G1 mice (Supplemental Figure 2, R-T). Since activation and subset expansion were best demonstrable in CD8+T-cells, we focused subsequent phenotyping experiments on CD8+T-cells.

### Variant APOL1 CD8+ T-cells show increased proliferation, central memory (CM) expansion, and cytokine expression following T-cell receptor stimulation, which is reversed by an APOL1 inhibitor

In PBMCs from healthy humans, we previously reported markedly increased *APOL1*-expression after TCR-stimulation of CD8+T-cells vs naïve T-cells (17). To study T-cell function with APOL1-RVs, we isolated CD8+T-cells from G0, and G1 mice (see methods) and stimulated them ex vivo with anti-CD3/CD28 (Supplemental Figure 3A) (24). We first evaluated *APOL1* expression in naïve and stimulated CD8+T-cells. APOL1 transcripts were similar in naive G0- and G1-CD8+T-cells, but proliferated G1-T-cells had significantly greater *APOL1* (Figure 3A). Consistent with this, total CD8+T-cells from G1-mice (with a greater proportion of CD8+CD44+T-cells) showed increased APOL1 vs G0-CD8+T-cells (Figure 3B). Ex vivo-stimulated G1-CD8+T-cells showed enhanced proliferation vs G0-CD8+T-cells (Figure 3, C and D). We isolated naïve T-cells from G1 and G0 spleens and stimulated them ex vivo (Supplemental Figure 3A). We confirmed the purity of isolated naïve CD8+T-cells (Supplemental Figure 3B). *Ex vivo* stimulated naïve G1- or G2-CD8+T-cells also showed significantly increased proliferation (Figure 3, E and F, Supplemental Figure 3, C and D) vs G0-CD8+ naïve T-cells, reporting a CD8+T-cell intrinsic phenotype. TCR-stimulated naïve G1-CD8+T-cells showed increased CD69+T-cells (recently activated), and increased CD44+T-cell proportions with expansion of TCMs. (Figure 3, G-I). This was accompanied by reduced viability and increased Annexin-V positivity in G1-CD8+T-cells (Supplemental Figure 3, E-H). Altered proliferation, activation, expansion, viability, and apoptosis in G1-CD8+T-cells were mitigated by MZ-302, an APOL1-inhibitor (Figure 3, G-K, Supplemental Figure 3, E-H) (25). MZ-302 had no effect on activation, or proliferation of WT-B6 CD8+T-cells without *APOL1* (Supplemental Figure 3, I and J).

**Figure 3.**
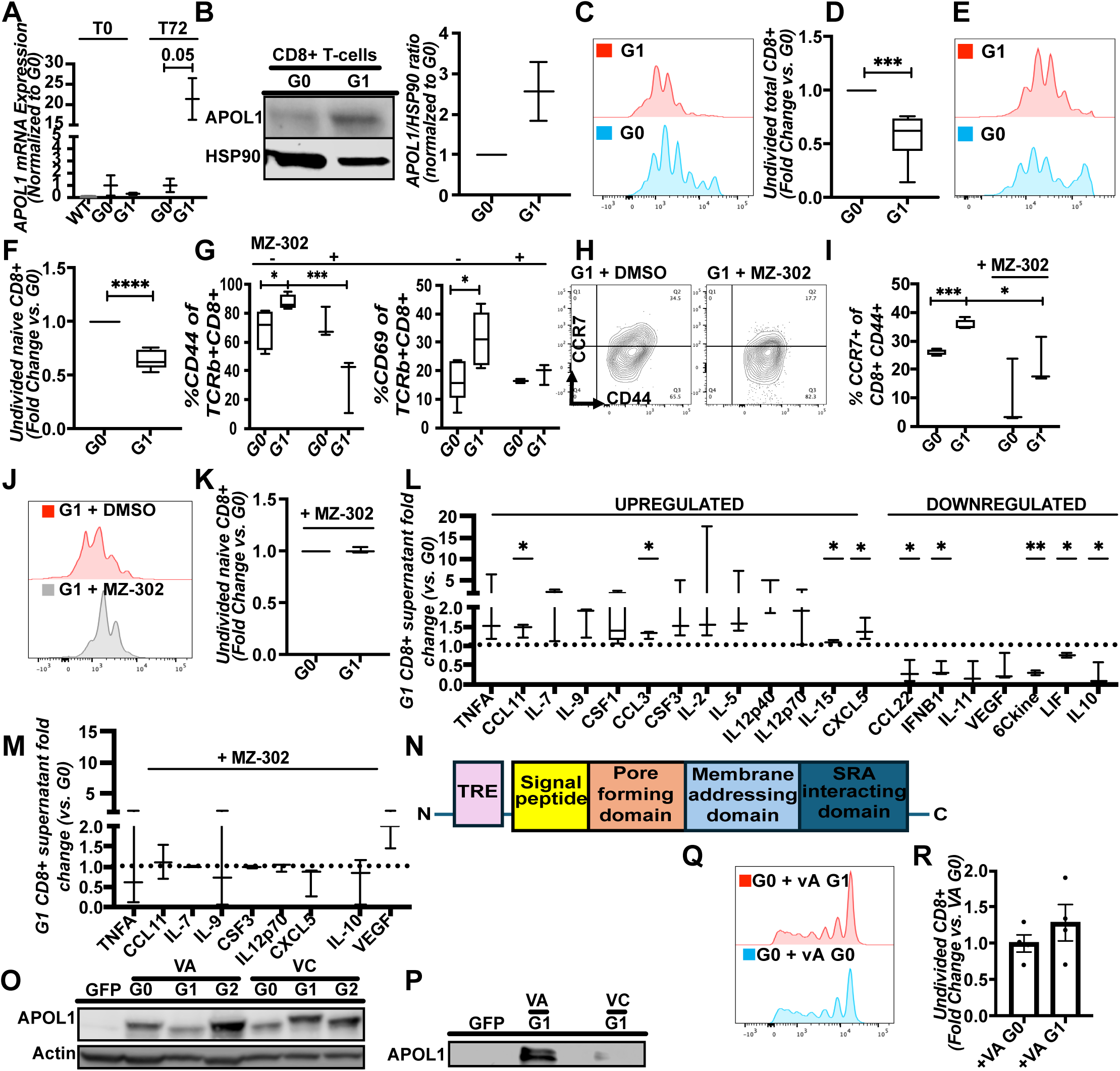
G1-CD8+T-cells show increased proliferation/cytokine expression with TCR stimulation, reversed by APOL1 inhibition. (A) *APOL1* mRNA in naïve G0-, G1-CD8+T-cells at baseline (T0) and post-TCR stimulation (T72). (B) Representative APOL1 immunoblot and quantification from G0-, G1-CD8+T-cell lysates (n = 10 mice). (C) Proliferation plots and (D) quantification (fold change) of undivided total G0-, G1-CD8+T-cells stimulated with anti-CD3/CD28. (E) Proliferation plots, and (F) quantification of undivided naive G0-, G1-CD8+T-cells stimulated with anti-CD3/CD28. (G-I) post-TCR-stimulation with/without MZ-302, (G) proportions of CD8+CD44+T-cells and CD8+CD69+T-cells, (H) representative flow plots and (I) proportions of CD8+CD44+CCR7+T-cells in G0-, G1-CD8+T-cells. (J) Proliferation plots, (K) quantification of undivided naive G0-, G1-CD8+T-cells post-MZ-302. (L-M) Box-Whisker plots show (L) G1-CD8+T-cell supernatant cytokine level fold changes (normalized to G0), (M) with MZ-302. (N) APOL1 protein/functional domains. Immunoblots from (O) cell lysates, (P) supernatant of HEK293-T-cells overexpressing APOL1 protein (VA-G0, -G1, -G2) or deletion constructs (VC-G0, -G1, -G2) probed for APOL1/β-Actin. (Q) Proliferation plots of G0-CD8+T-cells treated with VA-G1 or VA-G0 supernatant. [Bar graphs show fold changes of (R) undivided CD8+T-cells. Box-whisker plots show median and range of triplicates pooled from >5 mice each; bar graphs show mean ± SEM. Fold changes compared by paired T-test; *p<0.05, **p<0.01, ***p < 0.001, ****p<0.0001; MZ-302=APOL1 inhibitor]

To assess cytokine production, we performed cytokine profiling of supernatants from ex-vivo stimulated G1-, and G0-CD8+T-cells. G1-T-cell supernatant demonstrated upregulation of 13/44 cytokines while 7/44 were downregulated in three independent replicate experiments (N>5 mice/experiment; Supplemental Table 1). Pro-inflammatory cytokines canonically associated with TCR activation, such as TNFA and IL2, were upregulated while immunoregulatory cytokines were downregulated including IL10, VEGF, IFNB, IL11, and CCL22 (Figure 3L). Treatment with MZ-302 (10 uM) reversed both up- and down-regulation of cytokines in G1- vs G0-T-cell supernatant (Figure 3M).

APOL1 is the only secreted member among APOL proteins (26), and we next evaluated the role of secreted APOL1 in T-cell activation. We generated lentiviral, DOX-inducible, APOL1 overexpression constructs with either complete sequences (VA-G0, VA-G1), or with a deletion of the 21 amino-acid, signal peptide (VC-G0, VC-G1) (Figure 3N). In HEK293T-cells, inducible overexpression of these constructs was confirmed with immunoblots of cell lysates after 24 hours of overexpression (Figure 3O). Supernatant immunoblotting confirmed that only VA-, but not VC-construct expressing cells, had secreted APOL1 protein (Figure 3P). We next proliferated G0-T-cells ex vivo (anti-CD3/CD28) in the presence of either VA-G1- or VA-G0-APOL1 supernatant from overexpressing HEK293T-cells. Proliferation and viability of G0-T-cells were unchanged by the addition of secreted G1- or G0-APOL1 (Figure 3, Q and R, Supplemental Figure 4, A-C). Furthermore, induction of *APOL1* (with DOX), showed reduced viability in VA-G1 or VC-G1-lines vs corresponding G0-lines (Supplemental Figure 4D). Together, these data demonstrate a role for variant APOL1 in T-cell proliferation and cytokine expression, and that secreted variant-APOL1 plays a minimal role in these phenotypes.

### Variant BAC-Tg mice phenocopy cytokine excess observed in APOL1-RV patient sera following viral stimulation

To study immune responses to viral stimuli in G1-BAC-Tg mice, we injected the viral nucleic acid mimic Poly(I:C) intravenously into G0- and G1-BAC-Tg mice then profiled serum cytokines (on days-1, -7) and T-cell responses (day-7) (Figure 4A). On day-7, total splenocytes as well as adaptive and innate immune cell proportions were similar between G0- and G1-spleens (Supplemental Figure 5, A-F). G0-spleens showed increased CD4+CD44+ and CD8+CD44+T-cells vs untreated G0-mice; however, G1-spleens showed further increases in these cell proportions (Figure 4B). Poly(I:C)-treated G1-spleens also had lower Treg proportions than G0 (Supplemental Figure 5G). Multiplex cytokine profiling of sera from G1 mice again showed upregulation of pro-inflammatory cytokines starting from day-1, which persisted and significantly increased by day-7 (Figure 4C; 11/45 increased in G1 vs 0/45 significantly increased in G0). IFNG was below the detection threshold in the multiplex assay but was increased in G1-serum by ELISA at day-7 (Figure 4D). Since serum reflects both T-cell- and non-T-cell-cytokines, we specifically evaluated CD8+T-cell cytokine expression in response to ex-vivo treatment of splenocytes with Poly(I:C). G1-CD8+T-cells showed increased TNFa, IL-1b, IL-2, and IFNG expression with ex vivo Poly(I:C) treatment (Figure 4, E-H).

**Figure 4.**
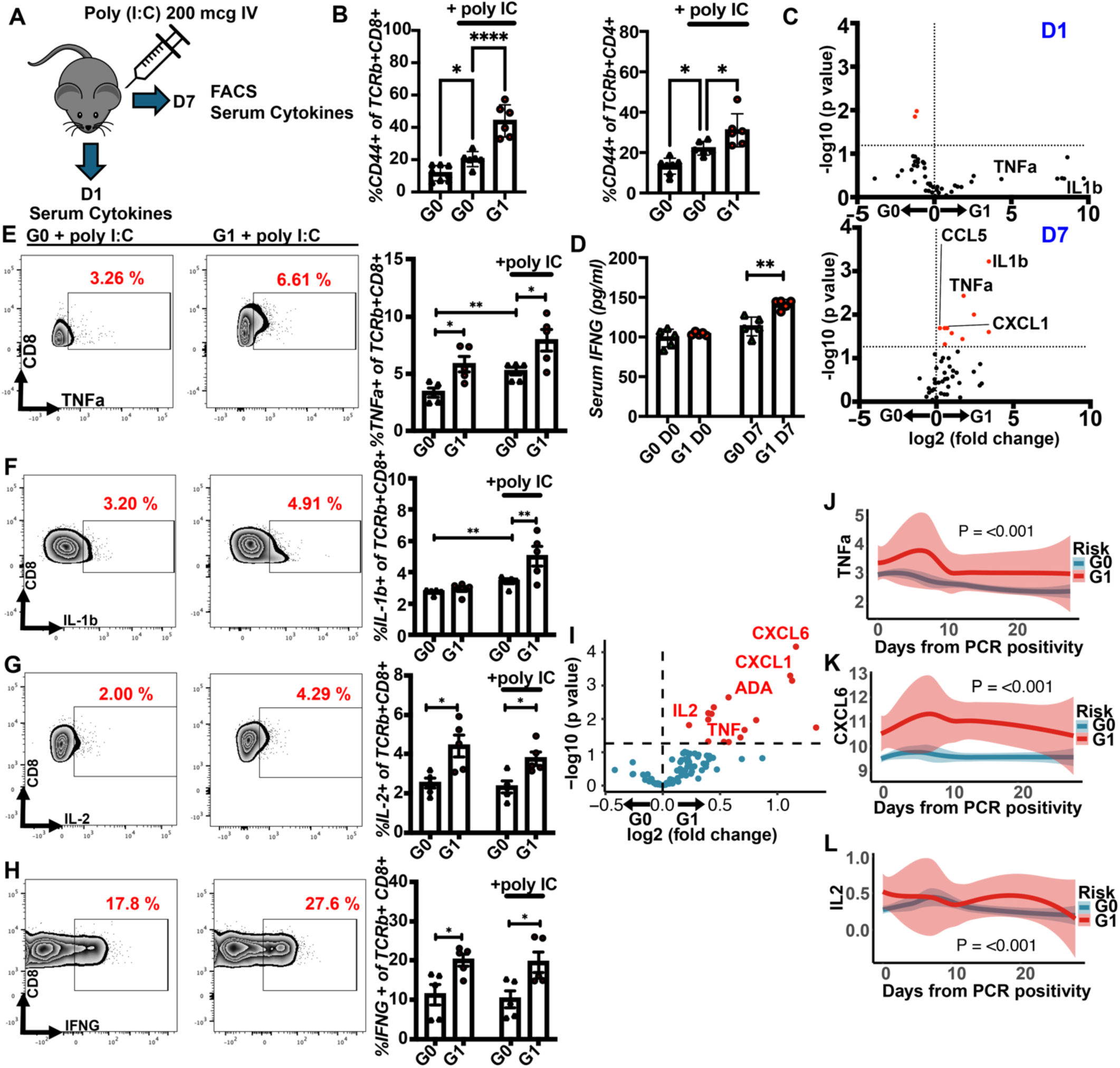
CD8+T-cells and sera from variant BAC-Tg mice phenocopy cytokine excess observed in APOL1-RV patient sera following viral stimulation. (A) Schema of *in vivo* Poly(I:C) study. (B) Proportions of CD8+CD44+ and CD4+CD44+T-cells with/without Poly(I:C) treatment. (C) Volcano plots of multiplex serum cytokine profiles at day 1 and at day 7 following Poly(I:C) treatment, a horizontal dotted line here corresponds to P-value =0.05 (unpaired T-test). (D) Serum IFNG levels by ELISA in G1 and G0 at day 0 and after 7 days from Poly(I:C) treatment. (E-H) Representative flow plots and bar graphs depicting pro-inflammatory cytokine production by CD8+T-cells- (E) TNFa, (F) IL-1b, (G) IL-2, and (H) IFNG in G1 vs G0 following an *ex vivo* Poly(I:C) treatment. (I) Volcano plot of relative cytokine expression changes during COVID-19 in G1 vs G0 individuals in CHARM study. (J-L) Line graphs of serum levels of (J) TNFa, (K) CXCL6, and (L) IL-2 (X-axis shows time day 0 to 28 from PCR positivity; Y-axis shows normalized protein expression by OLINK). The bar graphs display mean ± SEM. *p<0.05, **p< 0.01, ****p<0.0001, circles/triangles represent individual mice.

A viral infection classically associated with systemic cytokine excess and APOL1-induced disease is COVID-19 (27) (28). To understand whether cytokine excess observed in BAC-Tg-G1 mice is also identifiable in APOL1-RV patients who were infected with COVID-19, we used data from the CHARM-study where previously healthy US marines had serial sampling of sera before, during, and after SARS-CoV-2 infection in ambulatory settings. These marines remained outpatient during infection and were minimally symptomatic. We resolved the APOL1 genotype using RNA-seq data as described before (17). Table 1 lists demographics of patients used in this study. We analyzed serum cytokine trends from O-link data (96-plex) using linear mixed models that allowed us to incorporate multiple time points within each marine (see methods). G1-individuals showed significantly higher levels of 35/96 cytokines (P<0.05) accounting for multiple values obtained from each individual (any G1 vs G0/G0, n=6 vs 50) (Figure 4I, Supplemental Table 2). Similarly, 20/96 cytokines were significantly elevated in marines with any variant (i.e. any G1- or G2; n=12) vs G0/G0 (n= 50) (Supplemental Figure 5H, Supplemental Table 3). APOL1-RV marine sera also showed persistence of elevated cytokine levels (Figure 4, J-L and Supplemental Figure 5, I-K). Supplemental Figure 5L depicts differentially detected cytokines in COVID sera that overlapped with upregulated cytokines in CD8+T-cell supernatants, notably IL2 and TNFa. These data show increased T-cell activation and cytokine responses with APOL1-RV following viral stimulation, and relay human relevance for these findings.

**Table 1.** Demographic information of the marine COVID cohort (CHARM-study)(57), between *APOL1* genotypes.

| Participant *APOL1* genotype | All (n = 75) | G0 (n = 63) | G1<sup>a</sup> (n = 6) | G2<sup>b</sup> (n = 6) | P value |
| --- | --- | --- | --- | --- | --- |
| Male (%) | 64 (85.3) | 52 (82.5) | 6 (100) | 6 (100) | 0.51 |
| Age (y), Mean ± SD | 19.55 ± 2 .06 | 19.51 ± 2.14 | 19.83 ± 1.72 | 19.67 ± 1.75 | 0.93 |
| Race<sup>d</sup>, n (%) |  |  |  |  | **< 0.001** |
| EUR | 26 (34.7) | 26 (41.3) | 0 (0) | 0 (0) |  |
| AA | 10 (13.3) | 3 (4.8) | 5 (83.3) | 2 (33.3) |  |
| Hispanic | 14 (18.7) | 12 (19) | 1 (16.7) | 1 (16.7) |  |
| Other | 8 (10.7) | 5 (7.9) | 0 (0) | 3 (50) |  |
| Unknown | 17 (22.7) | 17 (27) | 0 (0) | 0 (0) |  |
| Symptoms | 35 (46.7) | 32 (50.8) | 2 (33.3) | 1 (16.7) | 0.29 |

a: Including WT/G1, G1/G1 and G1/G2.
b: Including WT/G2 and G2/G2.
c: Categorical variables were tested with chi-square and numeric variables with one-way ANOVA.
d: Self-reported race.
e: The symptoms are all mild according to the original paper(57).

### Variant APOL1-BAC-Tg mice demonstrate accelerated allograft rejection in an allogenic heart transplant model

Previous data suggest a role for APOL1-RVs in kidney allograft rejection (17-19). To evaluate the effect of APOL1-RVs in an allotransplantation model, we generated bone marrow chimeric mice from G0- and G1-BAC-Tg (in a B6/45.2 background) in B6/45.1 hosts. B6/45.2 chimerism was confirmed at 4 weeks in B6/45.1 hosts (both H-2^b^) (Supplemental Figure 6A). We performed mismatched heterotopic heart transplants where BALBc/J (H-2^d^) donor hearts were transplanted into G0- or G1-chimeric mice (n=4 each) with co-stimulation blockade using CTLA4-Ig (Figure 5A). Mice were monitored for rejection until 28 days (see methods and Supplemental Figure 6B). At termination, surviving allografts (n=3 each) were homogenized and the infiltrate examined by FACS. Hematoxylin and eosin staining of allografts revealed increased immune infiltrates and tissue necrosis in allografts of G1-recipients (Figure 5B). FACS analysis further showed increased immune infiltrates in allografts from G1-chimeric recipients including increased CD45+ cells (Figure 5C) and total T-cells (Figure 5D). CD8+T-cells as well as CD8+CD44+T-cells were both significantly increased (Figure 5, E and F). CD4+T-cells, CD4+CD44+T-cells, and innate immune cells (CD3-B220-) also tended to be increased in G1-chimeric recipient hearts (Supplemental Figure 6, C-E).

**Figure 5.**
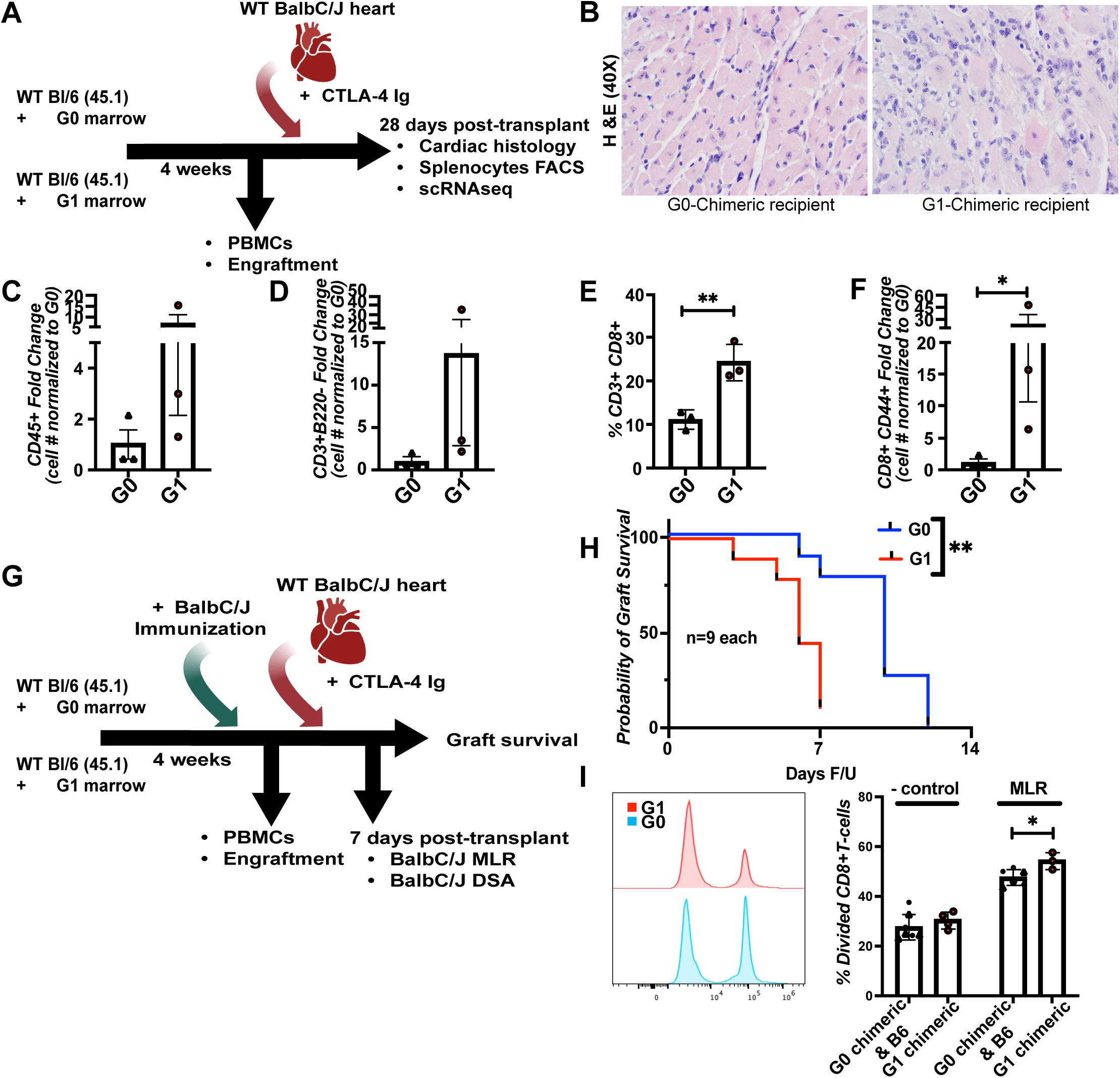
Variant APOL1-BAC-Tg mice demonstrate accelerated allograft rejection in an allogenic heart transplant model. (A) Schema of the allogenic heart transplant model. G1 or G0 bone marrow (B6/45.2) were transplanted into B6/45.1 syngeneic hosts to create G1- or G0-chimeric mice. Heterotopic allogenic transplants were performed with BALBc donor hearts into G1- or G0-chimeric recipients under costimulation blockade (n=4 each). (B) Hematoxylin and eosin photomicrographs of the allograft show excessive immune infiltrates in G1-chimeric recipients vs G0. (C-F) FACS analysis of the allograft for (C) CD45+CD45.2+cells, (D) total T-cells (CD3+B220-), (E) CD8+T-cells, and (F) CD8+CD44+Tcells (n=3 each). (G-I) G0- or G1-chimeric mice were sensitized with BALBc splenocytes (5×10^6^), followed by BALBc-heart transplantation (n=9 each). (H) Kaplan-Meier curves show allograft survival data in G1-chimeric recipients vs G0 controls in this sensitized model. (I) Proliferation plot and bar graphs show a mixed lymphocyte reaction from CD8+T-cells (to stimulator BALBc splenocytes) obtained at day 7 from a subset of transplants, comparing G1-chimeric recipients- with G0-chimeric- and WTB6 non-chimeric recipients (solid black circles) of BALBc hearts. Bar graphs display mean ± SEM. *p <0.05, **p<0.01.

Next, we immunized G0- and G1-chimeras with BALBc-splenocytes (5×10^6^ cells/mouse) followed by BALBc/J-heart transplantation with co-stimulation blockade (n=9 each, sensitized model) (Figure 5G). Here, BALBc-immunized G1-chimeric recipients showed significantly reduced allograft survival compared to G0-chimeric recipients (Figure 5H). Splenocytes from a subset of immunized and transplanted G1-chimeric mice at day-7 post-transplantation showed higher CD8+ and CD4+T-cell activation vs G0-chimeras (Supplemental Figure 6, F and G]. At day-7, a mixed lymphocyte reaction using BALBc splenocytes as stimulators showed increased proliferation of CD8+T-cells from G1-chimeric recipient spleens vs G0-chimeric recipients or non-irradiated, non-chimeric WT-B6/45.1 recipients (Figure 5I). Hence, chimeric G1-BAC-Tg mice demonstrated enhanced CD8+T-cell graft infiltration, antigen-specific responsiveness and accelerated rejection vs G0-chimeric recipients in an allogenic heart transplant model, analogous to observational data from human transplant cohorts.

### Transcriptome analyses from BAC-Tg CD8+ T-cells relays a role for enhanced Calcium-mediated Calcineurin/NFAT signaling in variant mouse T-cell activation and expansion

To investigate mechanism of phenotypic changes in APOL1-RV T-cells, we performed mRNA sequencing of ex-vivo stimulated naïve CD8+T-cells from G1 and G0 mice (n=3 vs 4, respectively; see methods). Batch normalized principal component analyses of transcriptome-wide profiles showed clustering of CD8+T-cells by genotype (PC1 explained 32%, and PC2 28% of variance; Figure 6A). Significantly differentially expressed genes (DEGS) (445 upregulated and 300 downregulated; Limma P<0.05; Figure 6B, Supplemental Table 4) were identified, and ranked for enrichment analyses. *APOL1* transcript was increased in G1-T-cell transcriptome despite similar genomic copy numbers, consistent with QPCR data (Figure 3A vs Supplemental Figure 7A). Top enriched pathways at different P-value thresholds are plotted in Figure 6C. Canonical immune response pathways related to TCR-stimulation were identified as significantly enriched in G1-CD8+T-cells including IL2/Stat5-, NFKB-, and Calcineurin/NFAT-signaling. Interestingly, multiple pathways related to calcium transport/calcium signaling (Figure 6C, Supplemental Figure 7, B and C) were identified as enriched. Selected dysregulated genes representing calcium channels or calcium-activated signals are labelled in Figure 6B.

**Figure 6.**
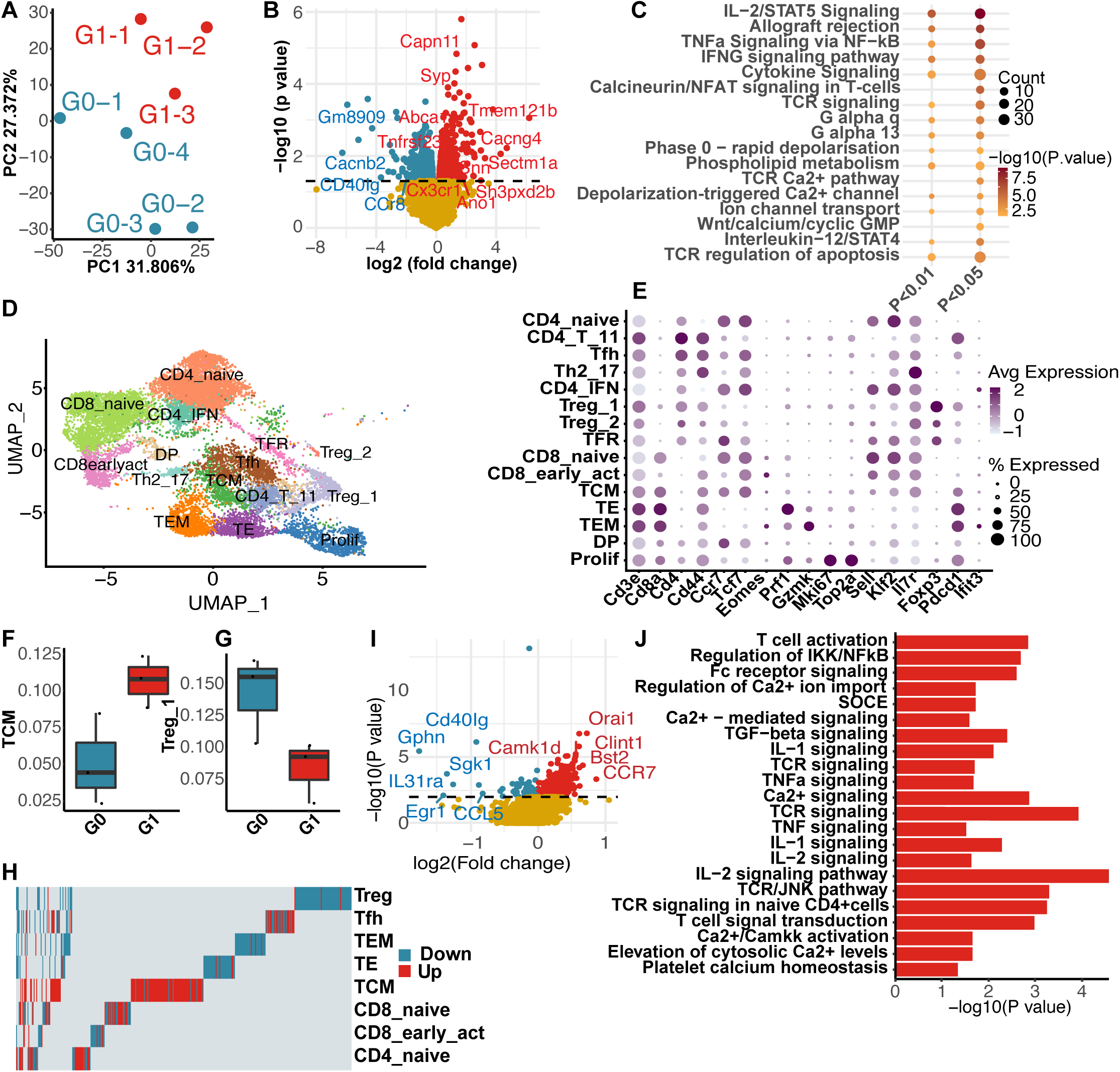
Transcriptome analyses relays enhanced Calcium-mediated Calcineurin/NFAT signaling in G1-CD8+T-cells. (A) PCA analysis of CD8+T-cell transcriptome of G1- vs G0-BAC-Tg. (B) Volcano plot of significant DEGs in G1 vs G0 transcriptomes. (Dashed line P=0.05; Selected Ca^2+^ signaling related transcripts are highlighted) (C) Functional enrichment analyses of significant DEGs (at P<0.01 and P<0.05, respectively). (D) UMAP of ScRNAseq of CD3+T-cells from G1- and G0-recipient allografts (n=3 each). (E) Bubble plot shows expression of canonical T-cell markers for subset annotation. Boxplots show proportions of infiltrating (F) TCMs and (G) Tregs in each sample (thick bar=median; box=25^th^-to-75th percentile, whiskers reach observations within 1.5 box-heights). (H) Significant DEGs (P<0.01) within each T-cell subset in G1 vs G0-comparisons. (Each column represents one gene, red = up-regulated, blue = down-regulated) (I) Volcano plot of the DEGs from infiltrating TCMs in G1 vs G0 comparisons (Dashed line P=0.01). (J) Functional enrichment analyses with top 50 DEGs (ranked after downsampling) in TCMs. IFN: interferon, CD8_early_act: activated CD8 cells in early phase, DP: CD4/CD8 double positive, TEM: effector memory, TE: effector T-cells, TCM: central memory, Tfh: T-follicular helper, TFR: T-follicular regulatory cells, Treg: Regulatory T-cells, Prolif: proliferating cells.

To study G1-CD8+T-cell subsets, we evaluated the single-cell transcriptome of graft infiltrating T-cells from the BALBc/J cardiac allografts harvested from chimeric G1- or -G0 recipients at 4-weeks (N=3 each; (Figure 5A)). We flow-sorted live CD3+ cells from allografts and performed single-cell RNA sequencing using a hashtagging approach (29) to multiplex the 6 samples during sequencing (n∼17000 total cells), (Supplemental Figure 7, D and E). As also seen in Figure 5E, deconvolution of hashtags revealed that significantly more T-cells had infiltrated allografts in G1-recipients (Supplemental Figure 7F). T-cell subsets were annotated as described before (30, 31), and 22 clusters were identified (Figure 6, D and E). Total numbers of all CD8+T-cell subsets were significantly increased in G1-chimeric recipients (Supplemental Figure 7F). G1-Sample #5 was an outlier in these analyses with high numbers of naïve CD4+ and CD8+T-cells. Hence, we evaluated the proportions of infiltrating cells within each graft after excluding naïve T-cells (Supplemental Figure 7G). We observed that the proportions of TCMs, T-effectors (TE), and T-effector/memory (TEMs) were all increased in G1, whereas proliferating- and T-reg proportions were reduced in G1- vs G0-chimeric recipients (Figure 6, F and G, Supplemental Figure 7H).

We then evaluated DEGs within each of these annotated cell clusters. Due to the imbalance in the number of infiltrating cells between G1 and G0 (Supplemental Figure 7F and Supplemental Figure 8A), we performed random down-sampling (bootstrapping). This approach equalized the number of cells analyzed in each iteration between G0- and G1-recipients, minimizing the potential for bias during DEG identification. Within each cell type, significant DEGs were identified from every bootstrapped test and ranked by the frequency of their occurrence. DEGs identified from ≥50% of bootstrapped samples were considered as robust DEGs and then ranked by their mean fold change. Supplemental Table 5 lists top-ranked DEGs (by average fold change) identified in each cell type. Enrichment analyses performed from DEGs in TEs (T effectors), TEMs (T effector memory), Naïve-CD8, and proliferating T-cells by the down-sampling approach are shown in Supplemental Figure 8B. G1-Naïve-CD8+T-cells showed enrichment of canonical TCR-signaling, IL2/Stat5- and TNF/NFKB signaling pathways suggesting basal activation in naïve G1-T-cells (Supplemental Figure 8B). TCMs demonstrated the greatest transcriptional perturbation by DEG number (n=642; Figure 6, H and I). In TCMs, the top 50 significant DEGs (by fold change) were subjected to enrichment analyses (Figure 6J). Consistent with bulk seq data from TCR-activated CD8+T-cells, Calcium-Calmodulin-kinase axis, canonical TCR activation with calcineurin NFAT activation, Calcium signaling and TNF/NFKB signaling were significantly enriched in G1-TCM cells. Interestingly, the top-ranked upregulated DEG in G1-TCMs was CCR7, consistent with FACS data of G1- BAC-Tg mice. ORAI1, a calcium channel activated by STIM1 with a key role in store-operated calcium entry (SOCE), was highly upregulated in G1-TCMs (Figure 6I). As sensitivity analyses, we excluded G1-Sample #5 and repeated the down-sampling approach to re-identify significant DEGs. DEGS from this sensitivity analyses demonstrated significant overlap with originally identified DEGs within each cell type including TCMs, naïve-CD8+ and naïve-CD4+T-cells (Hypergeometric P<0.001; Supplemental Figure 8C). Similarly, DEGS identified with or without the down sampling approach also showed significant overlap (Supplemental Figure 8C), confirming our results.

Since IL-1b was increased by Poly(I:C), and enriched in TCM transcriptome (Figure 6J), and the NLRP3 inflammasome was implicated in APOL1-disease (9), we tested this pathway in G1-CD8+T-cells by stimulating splenocytes *ex vivo* using Poly(I:C) with/without MCC950 (inflammasome inhibitor) (32). Here, we observed no inhibition of TNFa, IL2, and IFNG production from G1-CD8+T-cells treated with MCC950, although IL-1b excess was inhibited (Supplemental Figure 8, D-G), suggesting that the NLRP3 inflammasome does not play the major role in APOL1-G1-CD8+T-cell cytokine excess.

### Role of ER calcium depletion and SOCE in T-cell activation by variant-APOL1

Our unbiased bulk- as well as single-cell-transcriptomics relayed enrichment of calcium (Ca^2+^) facilitated signaling pathways in G1-CD8+T-cells. APOL1 is a monovalent cation channel (33) partly localized in the plasma membrane (12, 13), and it was recently reported that increased channel activity in APOL1-RVs at the cell surface (vs G0-APOL1) initiated a mechanism for cytotoxicity in epithelial cells by altering the resting membrane potential (11, 14, 15). Here, plasma membrane depolarization led to phospholipase C activation via GPCRs with IP3-mediated Ca^2+^ channel activation (pathways also enriched in G1-CD8+T-cell transcriptomes (Figure 6, C-J), causing ER-Ca^2+^ leak and depletion (13). Importantly in T-cells, canonical TCR-activation impinges on IP3-mediated Ca^2+^ release leading to ER-Ca^2+^ depletion, in turn stimulating SOCE (via Calcium release-activated calcium- (CRAC) Orai-Stim1 channels) and Calcineurin activation (34). Indeed, *Orai1*, a key participant in SOCE, was highly upregulated in the G1-TCM transcriptome. We also confirmed that APOL1 expression occurred on the plasma membrane of CD8+T-cells as reported in epithelial cells (Supplemental Figure 9A).

To first investigate ER-Ca^2+^ stores, CD8+T-cells from G0- and G1-mice were isolated and stimulated *ex-vivo* for 48 hours (in plates coated with anti-CD3 for adhesion; n=5 mice each). ER-Ca^2+^ stores reflect a balance of Ca^2+^ entering the ER via sarco/endoplasmic reticulum calcium ATPase (SERCA) pumps (35), and Ca^2+^ release via the IP3R and ryanodine receptors (36). We assayed changes in cytosolic Ca^2+^ concentration ([Ca^2+^]_cyt_), which reflect efflux from ER-Ca^2+^ stores, by live cell imaging using Fluo-4-AM dye immediately after addition of Thapsigargin (a SERCA-inhibitor), and simultaneously precluding ECF-Ca^2+^ entry using EGTA. Here, both amplitude (Supplemental Figure 9B) and area-under-the- curve (AUC; (Figure 7, A-C)) of Thapsigargin-induced [Ca^2+^]_cyt_ peak were significantly reduced in the G1-CD8+T-cells, confirming ER-Ca^2+^ depletion in G1- vs G0-CD8+T-cells.

**Figure 7.**
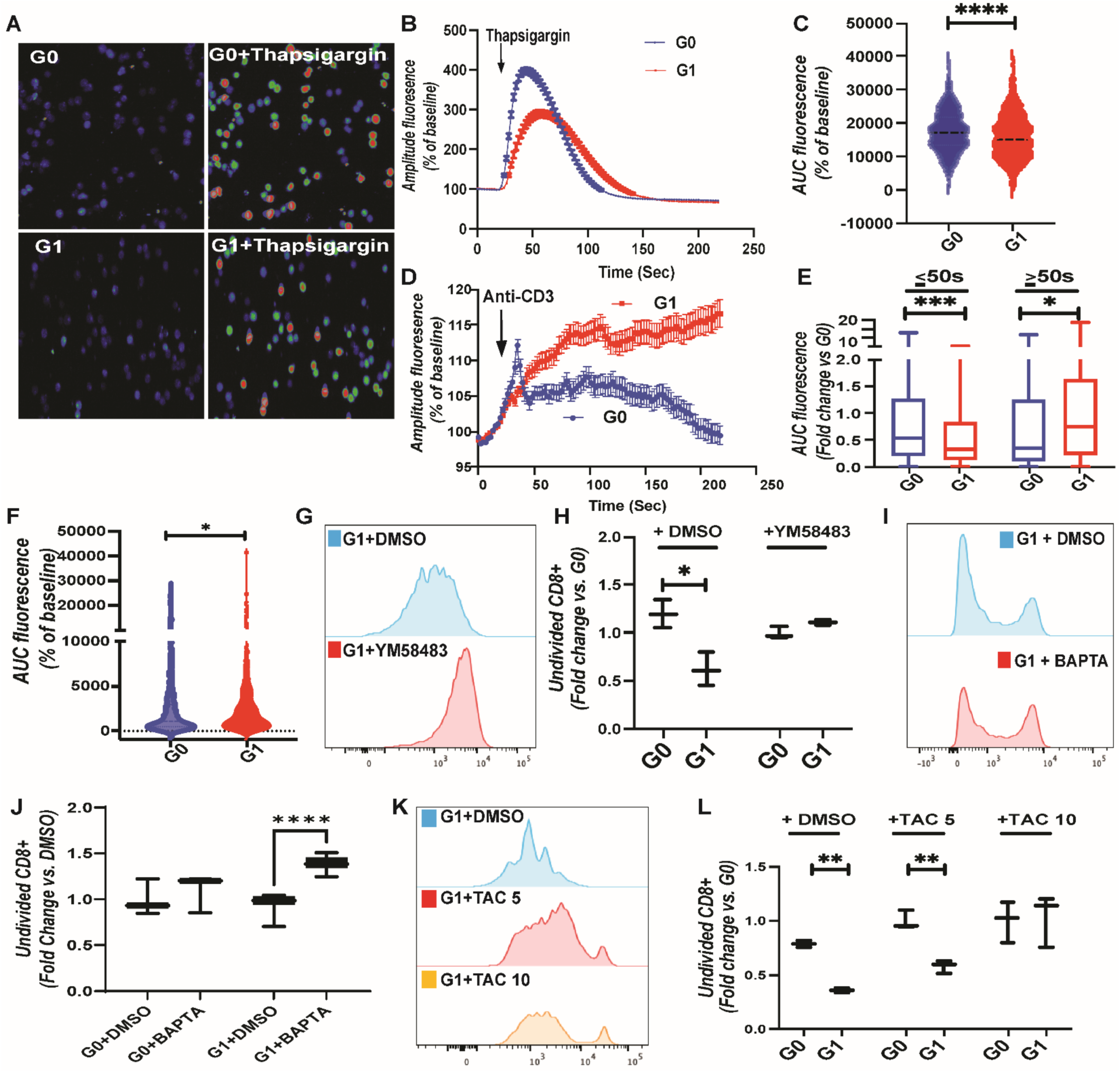
ER calcium depletion causes activation of variant APOL1 CD8+T-cells. (A-C) (A) Representative images- (dots = T-cells), (B) mean calcium flux curves-, and (C) corresponding average AUCs- of fluorescence measured from live G0- and G1-CD8+T-cells in response to Thapsigargin [N=5 mice each]. (D-F) (D) summarized calcium flux curve, (E) AUCs- of early vs late fluorescence (< or >50 seconds), and (F) -total fluorescence from CD8+T-cells from G1 vs G0 after anti-CD3-stimulation [(>500 cells, n>6 mice] (G) Representative plot and (H) quantified undivided total CD8+ T-cell of G1 vs G0 (fold change from triplicates of 5 mice) with YM-58483 at 50 nM. (I) Proliferation plot of G1-CD8+T-cells treated with BAPTA-AM (0.3 uM) and DMSO. (J) Quantified effect of BAPTA-AM (0.3 uM) on naïve CD8+T-cell proliferation (expressed as fold-change of undivided cells in G0- and G1-DMSO treatment (triplicates of 5 mice), respectively. (K) Representative plot of G1-CD8+T-cell treated with TAC at 5, 10 ng/mL (L) Effect of TAC at 5, 10 ng/mL on the proliferation of G0- and G1-CD8+T-cell (expressed as fold change of undivided naïve CD8+T-cells from triplicates of 5 mice). Box-whisker plots show median and range of triplicates pooled from >5 mice each. *p<0.05, **p<0.01, ***p<0.001, ****p<0.0001. AUC: Area-under-the-curve. DMSO: Dimethyl sulfoxide, TAC: Tacrolimus.

We surmised that ER-Ca^2+^ depletion in G1-CD8+T-cells would promote earlier engagement of SOCE mechanism during TCR stimulation. To study Ca^2+^ flux following TCR stimulation, CD8+T-cells from G0- and G1- mice (n=6 each) were isolated and stimulated *ex-vivo* (as above) and [Ca^2+^]_cyt_ assayed after fresh replenishment of anti-CD3. An average [Ca^2+^]_cyt_ curve summarizing all measured G1- and G0-CD8+T-cells is shown in Figure 7D. Again, G1-CD8+T-cells demonstrated a blunted early increase in [Ca^2+^]_cyt_ giving rise to a significantly decreased amplitude and AUC of the early peak (<50 seconds), and indicating a reduced ER-Ca^2+^ release, vs G0 (Supplemental Figure 9C and Figure 7E, respectively). However, delayed [Ca^2+^]_cyt_ release (>50 seconds after anti-CD3), as well as overall [Ca^2+^]_cyt_ – expressed as AUC – were significantly higher in G1-CD8+T-cells vs G0 (Figure 7, D-F). These data confirm ER-Ca^2+^ depletion in G1-APOL1 CD8+T-cells with blunted early rise in [Ca^2+^]_cyt_ after TCR-stimulation but followed by sustained [Ca^2+^]_cyt_ in later phases of TCR stimulation.

SOCE mediated by the Orai1-Stim1 complex replenishes ER-Ca^2+^, contributing to the later part of the [Ca^2+^]_cyt_ curve and promoting T-cell activation via Calcineurin/NFAT. Consistent with the role played by SOCE specifically in G1-CD8+T-cells, reduced doses of CRAC-channel inhibitor YM-58483 [50 nM, IC_50_ = 330 nM (37)] obviated the increased proliferation and the reduced viability encountered in ex vivo TCR-stimulated G1-CD8+T-cells vs G0 (Figure 7, G and H, Supplemental Figure 9D).

To further test the role played by [Ca^2+^]_cyt_ levels in G1-CD8+T-cell phenotype, we stimulated G1- and G0-CD8+T-cells in the presence of Ca^2+^-chelator BAPTA-AM (38). At higher concentrations (3 uM, 48-hrs), BAPTA-AM promoted cell death in T-cells; we therefore used 10-fold reduced concentrations of BAPTA-AM (0.3 uM) (39). G1-CD8+T-cells were more sensitive to 0.3 uM BAPTA-AM with inhibited proliferation compared to G0-CD8+ T-cells (Figure 7, I and J). The calcineurin/NFAT pathway was enriched in G1-CD8+T-cells (Figure 6C), and we reasoned that greater activation of this pathway by SOCE in G1+CD8+T-cells would render G1-cells more refractory to inhibition by calcineurin inhibitors. We therefore tested CD8+T-cell activation in the presence of varying concentrations of the calcineurin inhibitor Tacrolimus (TAC at 0-,5- or 10 ng/ml). G1-CD8+T-cells showed greater proliferation vs G0-cells at 0- and 5-ng/ml, but differences in proliferation were abolished only at 10 ng/ml (Figure 7, K and L). Similarly, TCM expansion was dose-dependently decreased in G1-CD8+T-cells at 5- and 10-ng/ml but not obviated (vs G0) at 10 ng/ml (Supplemental Figure 9E).

### APOL1-RV kidney transplant recipients show increased rejection with higher TAC trough levels

Our studies above suggested that APOL1-G1 variant CD8+T-cells sustain ER-Ca^2+^ leak and depletion(13), followed by increased SOCE, Calcineurin/NFAT signaling and resistance to TAC inhibition, promoting TCM expansion and rejection. To evaluate this in an independent human kidney transplantation cohort, we utilized data and samples from the CTOT-19 study. From this NIAID-funded, placebo-controlled, randomized trial (40), which tested the impact of TNF-alpha blockade using infliximab on graft outcomes (n=225), we obtained DNA of kidney recipients. We performed genome-wide genotyping using a GSA.v3 SNP array with imputation (17) (41) and resolved APOL1-genotypes (methods). Among 210 CTOT-19 recipients with genotype data, 65 (30.9%) had APOL1-RV genotypes (49/65 had G1). The clinical characteristics of the genotyped CTOT-19 cohort are in Table 2. Of the 210 recipients, 81 (38.57%) were AAs and 73% of AAs had APOL1-RV genotype.

**Table 2.**
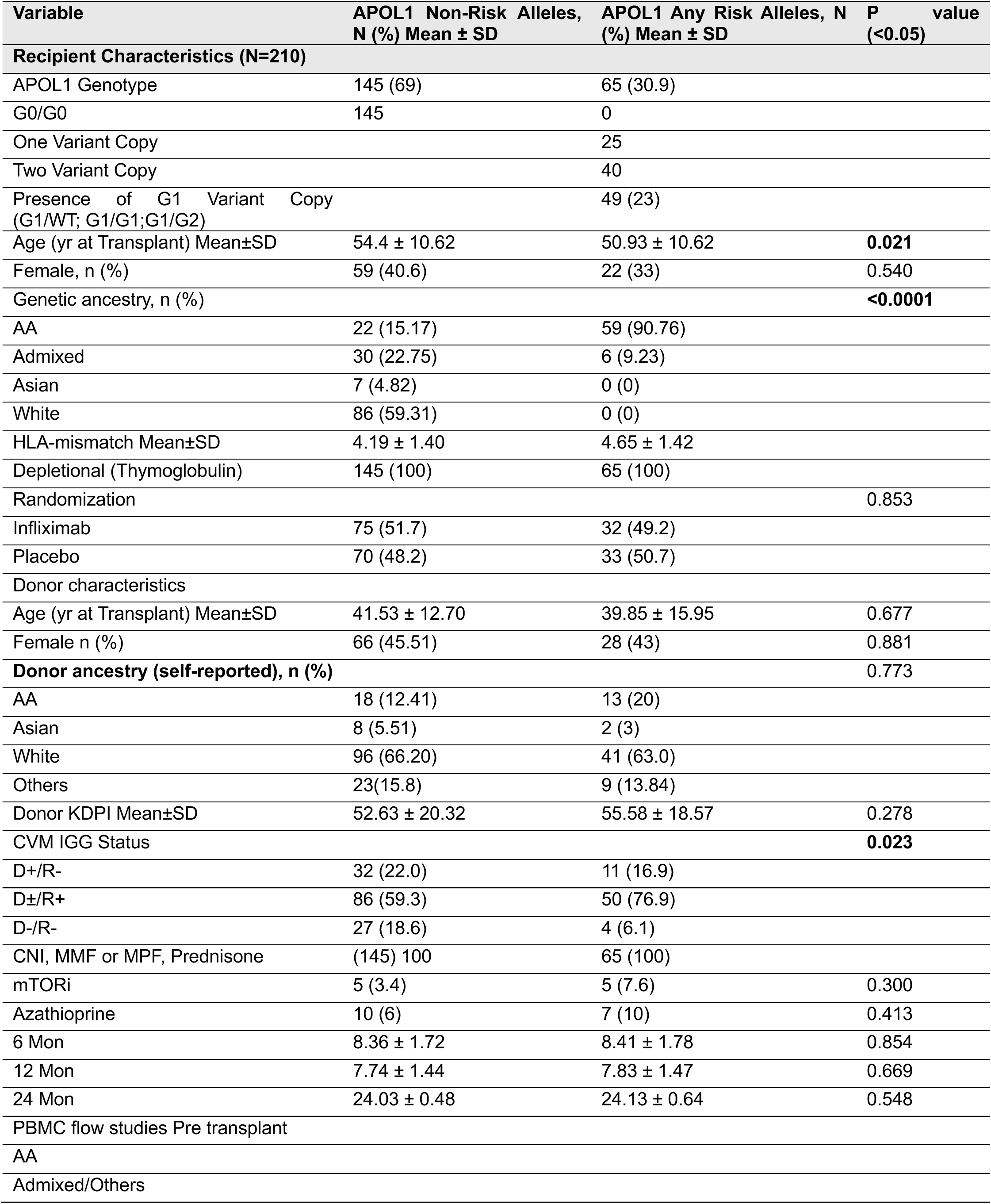
Clinical/demographic information of the genotyped CTOT19 cohort(40).

| Variable | APOL1 Non-Risk Alleles, N (%) Mean ± SD | APOL1 Any Risk Alleles, N (%) Mean ± SD | P value (<0.05) |
| --- | --- | --- | --- |
| **Recipient Characteristics (N=210)** |  |  |  |
| APOL1 Genotype | 145 (69) | 65 (30.9) |  |
| G0/G0 | 145 | 0 |  |
| One Variant Copy |  | 25 |  |
| Two Variant Copy |  | 40 |  |
| Presence of G1 Variant Copy (G1/WT; G1/G1;G1/G2) |  | 49 (23) |  |
| Age (yr at Transplant) Mean±SD | 54.4 ± 10.62 | 50.93 ± 10.62 | **0.021** |
| Female, n (%) | 59 (40.6) | 22 (33) | 0.540 |
| Genetic ancestry, n (%) |  |  | **<0.0001** |
| AA | 22 (15.17) | 59 (90.76) |  |
| Admixed | 30 (22.75) | 6 (9.23) |  |
| Asian | 7 (4.82) | 0 (0) |  |
| White | 86 (59.31) | 0 (0) |  |
| HLA-mismatch Mean±SD | 4.19 ± 1.40 | 4.65 ± 1.42 | |
| Depletional (Thymoglobulin) | 145 (100) | 65 (100) |  |
| Randomization |  |  | 0.853 |
| Infliximab | 75 (51.7) | 32 (49.2) |  |
| Placebo | 70 (48.2) | 33 (50.7) |  |
| **Donor characteristics** |  |  |  |
| Age (yr at Transplant) Mean±SD | 41.53 ± 12.70 | 39.85 ± 15.95 | 0.677 |
| Female n (%) | 66 (45.51) | 28 (43) | 0.881 |
| **Donor ancestry (self-reported), n (%)** |  |  | **0.773** |
| AA | 18 (12.41) | 13 (20) |  |
| Asian | 8 (5.51) | 2 (3) |  |
| White | 96 (66.20) | 41 (63.0) |  |
| Others | 23(15.8) | 9 (13.84) |  |
| Donor KDPI Mean±SD | 52.63 ± 20.32 | 55.58 ± 18.57 | 0.278 |
| **CVM IGG Status** |  |  | **0.023** |
| D+/R- | 32 (22.0) | 11 (16.9) |  |
| D±/R+ | 86 (59.3) | 50 (76.9) | |
| D-/R- | 27 (18.6) | 4 (6.1) |  |
| CNI, MMF or MPF, Prednisone | (145) 100 | 65 (100) |  |
| mTORi | 5 (3.4) | 5 (7.6) | 0.300 |
| Azathioprine | 10 (6) | 7 (10) | 0.413 |
| 6 Mon | 8.36 ± 1.72 | 8.41 ± 1.78 | 0.854 |
| 12 Mon | 7.74 ± 1.44 | 7.83 ± 1.47 | 0.669 |
| 24 Mon | 24.03 ± 0.48 | 24.13 ± 0.64 | 0.548 |
| **PBMC flow studies Pre transplant** |  |  |  |
| AA |  |  |  |
| Admixed/Others |  |  |  |

Rejection diagnoses in CTOT-19 were obtained from both local and central lab reports (Supplemental Table 6 and methods (40)). We included all rejection diagnoses made by central or local pathology (subclinical or clinical, ABMR or TCMR including borderline) as “AR” for this analysis. Among 210 patients, 34 patients developed AR by 6-months, 42 by 12 months, and 49 by 24-months (end of study). Within 6-, 12- and 24-months, patients with at least one biopsy who did not have AR diagnosis were considered as “no rejection” or “NAR”. We identified that APOL1-RV recipients had significantly increased incidence of AR by 6 months (Table 3). In time-to-event analyses, any APOL1-RV-, or APOL1-G1 -genotype, was associated with a significantly increased risk of AR by 6- and 12-months (Figure 8, A-D, respectively). The association between APOL1-RV copies and 6- or 12-month AR was identifiable in additive models (Supplemental Figure 10, A and B) (as also reported before (17, 19)). In bivariate Cox models for time to 6-month rejection, adjusted for recipient age, CMV status (i.e., covariates significantly different between APOL1-RV vs G0/G0 in Table 3), or randomization limb, APOL1-RV remained significantly associated with AR. Adjustment for genetic ancestry attenuated the significance of association of APOL1-RV with AR, albeit with similar hazard ratio (Supplemental Table 7). Including AR events up to 24-months tended to attenuate significance of association, suggesting a predominant association of APOL1-RVs with early AR (Supplemental Figure 10C).

**Figure 8.**
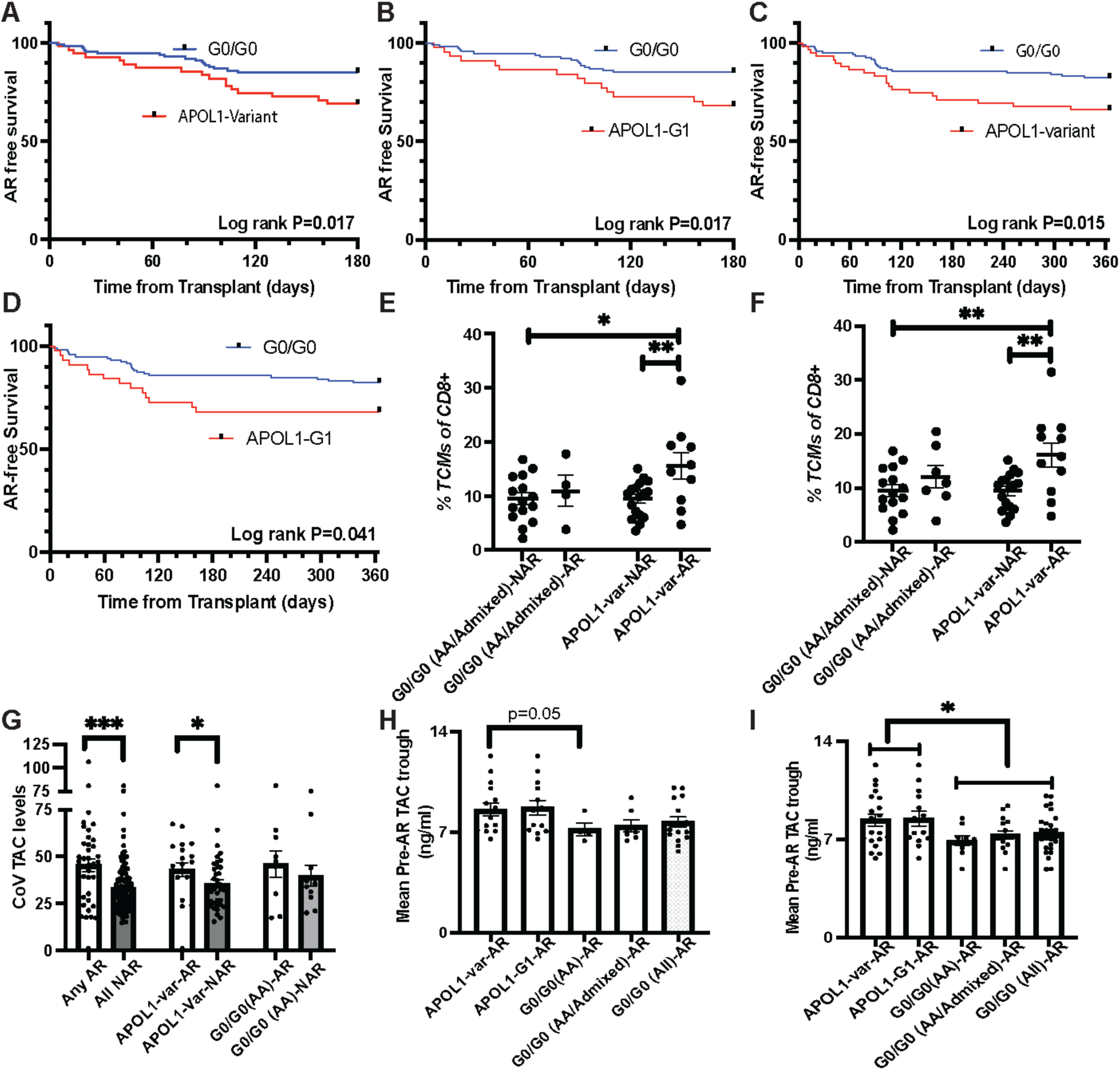
TCM expansion increases the risk of rejection in APOL1-RV kidney transplant recipients with higher TAC trough levels. CTOT-19 cohort was used to validate our findings. (A-D) Kaplan Meier curves show time-to-event analysis of (A) AR by 6-months in any APOL1-RV, (B) in APOL1-G1 recipients, and corresponding data for (C), (D) AR by 12-months, respectively, vs. G0/G0. (E-F) Dot plots of percentage TCMs (CD8+CD45RO+CD28+CD27+ among T-cells) pre-transplant by APOL1-genotype, (E) in AA/Admixed patients with/without AR by 6-months, and (F) in AA/Admixed patients with/without AR till 24-months. (G) The Coefficient of Variation (COV) of pre-AR TAC-troughs between AR and NAR, by APOL1 genotype. (H-I) Bar graphs show mean pre-rejection TAC-troughs by APOL1 genotype of (H) 6-month-AR vs NAR and (I) any time AR vs NAR [bar graphs display mean ± SEM.*p<0.05, **p<0.01].

**Table 3.** Acute rejection incidence according to APOL1 genotype.

| **Outcomes, N (%)** | **APOL1 Non-Risk Alleles, N (%)** | **APOL1 Any Risk Alleles, N (%)** | **P Value (<0.05)** |
| --- | --- | --- | --- |
| DGF (yes) | 46 (31) | 23 (35) | 0.635 |
| CMV Viremia (yes) | 21 (14.4) | 12 (18.4) | 0.539 |
| Presence Bk (yes) | 28 (19.31) | 19 (29.2) | 0.151 |
| HOSIM (yes) | 55 (37.9) | 31 (47.6) | 0.224 |
| ANYINFHD (yes) | 54 (37.2) | 31 (47.6) | 0.172 |
| Death 24 Mon (Yes) | 5 (3.4) | 2 (3.0) | 0.999 |
| **Rejection 6 Mon** |  |  |  |
| AR Central (yes)* | 4 (4.1) | 1 (2.7) | 0.999 |
| AR Local (yes)* | 13 (11.4) | 14 (25.4) | **0.025** |
| AR with Borderline Central (yes)* | 7 (7.2) | 5 (13.8) | 0.307 |
| Any AR (yes)* | 17 (14.91) | 17 (30.91) | **0.023** |
| **Rejection 24 Months** |  |  |  |
| AR Central (yes)* | 5 (4.9) | 3(7.5) | 0.686 |
| AR Local (yes)* | 21 (17.8) | 17 (28.8) | 0.119 |
| AR with Borderline Central (yes)* | 12(11.7) | 7 (17.5) | 0.414 |
| Any AR (yes)* | 29 (24.5) | 20 (33.9) | 0.214 |

*\*The presented percentages apply to biopsies for which results are available.*

We next investigated CD8+T-cell phenotype of APOL1-RV individuals who developed early AR (by 6-months; n=17) and compared these to APOL1-RV NARs of AA/Admixed ancestry (n=39) as well as G0/G0- AA/Admixed ancestry with AR (n=7), or with NAR (n=36; total 6-month AR data from n=99 (Table 3)). From FACS data collected at multiple time points, we focused on the pretransplant visit (visit 0), i.e., before initiation of immunosuppression or infliximab (Visit-0 FACS data available for 56/99 patients). Within this dataset, we compared pre-transplant CD8+T-cell phenotype of patients with/without AR by 6-months. We observed that APOL1-RV-ARs had significantly higher proportions of TCMs [CD8+CD45RO+CD28+CD27+T-cells, (Supplemental Figure 10D for gating strategy) than both APOL1-RV-NAR and G0/G0-NAR (Figure 8E), consistent with TCM expansion observed in murine data. Other CD8+ T-cell subsets including TEMRA [CD8+CD45RA+CD28-CD27-], T-effector [CD8+CD45RO+CD27-], and naïve CD8+ [CD8+CD45RA +CD28+CD27+] T-cell fractions were not significantly different between any of the groups (Supplemental Figure 9, E-G). The proportion of TCMs pre-transplant was still significantly higher in APOL1-RV-ARs when all AR episodes during the study were collated from APOL1-RV and G0/G0-AA/Admixed individuals (Figure 8F).

To investigate TAC dose dependency, we utilized mean TAC trough levels collected throughout the study. Overall, ARs had higher coefficients of variation (CoV) of TAC-troughs vs NARs, suggesting TAC variability and non-adherence as a predisposition for AR (Figure 8G). This finding was consistent among APOL1-RV recipients. Interestingly, APOL1-RV-ARs had higher TAC-trough levels before AR (i.e. pre-rejection TAC troughs) before 6-month AR (Figure 8H; P=0.05), and before any AR (Figure 8I), compared to APOL1-G0/G0-ARs (including both AAs/Admixed- and the entire-cohort), and suggest that AR occurred in APOL1-RV recipients despite higher TAC troughs. Together, these data validate the increased risk of AR with APOL1-RVs, and support CNI-mediated T-cell activation and TCM expansion seen in our murine studies (Summarized in Figure 9).

**Figure 9.**
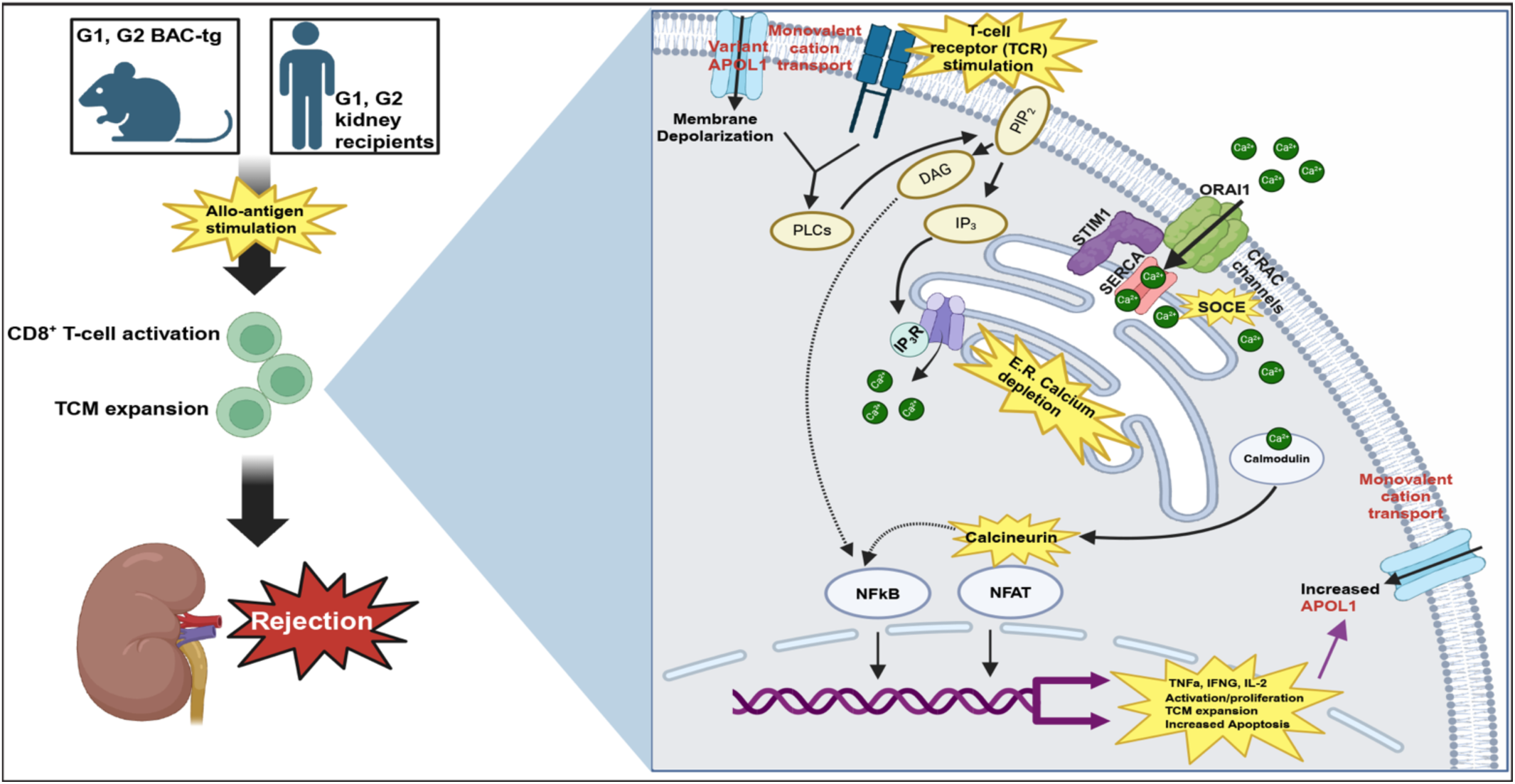
Summary Figure. Expression of APOL1-RVs (a cation transporter) in BAC-Tg CD8+T-cells causes membrane depolarization and activates PLCg-PIP_2_-IP_3_, driving sustained ER-Ca^2+^ leak via IP_3-_channels. Following TCR-ligation, pre-existing ER-Ca^2+^ depletion leads to sustained SOCE via STIM1-ORAI1. Downstream Ca^2+^-calmodulin-calcineurin activates NFAT and NFkB pathways, and enhanced cytokines, activation, and TCM-expansion, accompanied by activation-induced cell death. Clinically, recipients with APOL1-RVs developed increased AR when they had elevated TCMs pre-transplant and despite significantly higher tacrolimus levels vs. G0-AA-AR. Created in BioRender.

## DISCUSSION

APOL1-FSGS is consistent with a podocyte-centric paradigm of APOL1-RV toxicity, and data from kidney transplantation support increased DCAL in APOL1-RV donor kidneys vs G0/G0 (42). A growing body of clinical data have now also reported an association for APOL1-RVs in kidney recipients with TCMR and DCAL (17-19). It must be noted that epidemiologic reports cannot fully unravel the association of APOL1-RVs with rejection, since AA status, which is highly linked with APOL1-RV carriage (19, 43), is also associated with higher risk of rejection from clinical factors(20), pharmacogenomic loci influencing drug metabolism (21) and socio-economic indicators(19, 43). Thus, examining the role and mechanisms of APOL1-RVs in kidney rejection has considerable importance since an APOL1 inhibitor is already in phase-III human trials (44), and will have specific therapeutic implications. In the current series of experiments, we used a BAC-Tg mouse model to specifically investigate APOL1-G1, -G2 in T-cell responses leading to TCMR, and separate this from associations with AA-ancestry.

From these studies, we identified a clear excitatory CD8+T-cell phenotype in adult G1- and G2-BAC-Tg mice vs G0, with expansion of CD8+CD44+CCR7+T-cells (TCM) and a role in promoting inflammatory responses and allo-rejection. While we focused on backcrossed G1-mice for phenotyping experiments, we performed selected validation in G2-hybrid BAC-Tg mice, showing generalizability to both APOL1-RVs. The excitatory phenotype with APOL1-RV was observed in ex vivo stimulated naïve as well as antigen-experienced CD8+T-cells, and was ameliorated by an APOL1-antagonist, demonstrating a T-cell intrinsic, targetable phenotype in transplantation given the availability of APOL1-inhibitors.

Unbiased transcriptome analyses suggested altered Ca^2+^ signaling as an important underpinning mechanism in APOL1-RV-CD8+T-cells. ER-Ca^2+^ depletion has been previously implicated as a mechanism of cytotoxicity of APOL1-RVs in epithelial cells. In this model, plasma membrane depolarization by surface APOL1-RVs led to phospholipase C activation via GPCRs with persistent IP3-Ca^2+^ channel activation and ER-Ca^2+^ depletion (13). We experimentally confirmed surface expression of APOL1-RVs in CD8+T-cells and demonstrated depleted ER-Ca^2+^ stores in G1-CD8+T-cells. Furthermore, phospholipase C and GPCR-signaling were also consistently identified in G1-CD8+T-cell transcriptomes as dysregulated pathways supporting the involvement of this axis. Hence, our findings from G1-CD8+T-cells paralleled epithelial cells (12, 13) with surface APOL1-RVs causing secondary ER-Ca^2+^ depletion. However, *unlike in epithelial cells*, in T-cells, IP3-mediated ER-Ca^2+^ depletion drives SOCE-mediated [Ca^2+^]_cyt_ levels, and is central to Calcineurin/NFAT activation after TCR-stimulation. In line with this concept, we observed sustained increases in [Ca^2+^]_cyt_ levels with TCR-ligation in G1-CD8+T-cells, and perturbations of [Ca^2+^]_cyt_ with TCR-stimulation disproportionately impacted G1-CD8+T-cells. Inhibition of G1-CD8+T-cells proliferation required higher TAC levels, corresponding with increased Calcineurin/NFAT activation. Increased SOCE that we observed in G1-CD8+T-cells could also explain TCM expansion, as SOCE has been shown to be critical for upregulation of genes involved in oxidative phosphorylation promoting metabolic reprogramming and memory formation (45). Upregulated TNF in G1-CD8+T-cells could also be related to increased Ca^2+^-dependent or -independent NFKB activation downstream of TCR (46).

Our study has important implications. Our translational data suggests that recipients with APOL1-RVs may need higher levels of TAC to achieve sufficient immunosuppression. Notably, we observed that even at TAC levels of ∼10 ng/mL, there was incomplete inhibition of TCM expansion during TCR stimulation in G1-CD8+T-cells. Moreover, higher levels of TAC increase risks of infection/nephrotoxicity (47) and may not be clinically feasible. These provide therapeutic implications for our findings and the potential need for agents targeting APOL1 in APOL1-RV recipients, one of which is in clinical trials (44). Next, our data raises the possibility that individuals with APOL1-RVs who have a greater proportion of TCMs prior to transplantation may be at highest at risk for early AR and could be used as a risk-stratification tool for therapy or monitoring. Further studies, however, are needed to fully understand how TCM expansion modifies rejection risk among individuals with APOL1-RV. A similar role for recipient APOL1 in non-kidney solid organ rejection, also needs to be evaluated in future work. While the data presented here as well as other recently published studies suggest a role for recipient APOL1 genotype in kidney transplant outcomes, it is not yet clear whether the combination of recipient- plus donor- APOL1-RVs presents highest risk of graft loss, than either alone. Larger datasets with enrichment of donor- and recipient AAs with APOL1-RV genotype such as the APOLLO consortium will help clarify this question(48). Since viral insults, immune stimulation, and IFNG excess(10), all stereotypically induce APOL1-FSGS in native kidneys, a potential upstream role for T-cell activation in APOL1-mediated FSGS is also worthy of investigation.

Our data has some limitations. Haplotypic differences may influence APOL1-RV toxicity (49). All three BAC clones used here contain a lysine (K150) that is found only in the G0 (reference) haplotype. Our BAC-Tg mice therefore do not share the haplotype associated with APOL1-RVs in AAs. However, we support our murine data with multiple analogous findings in human cohorts which represent the correct AA haplotype. Hence, the phenotype in our variant mice result from the exonic changes in SRA-region induced by G1 and G2 mutations in APOL1 protein, and regardless of the background haplotype. We studied adult mice in this work and a role for APOL1-RVs during thymic development influencing adult T-cell phenotype was not addressed. Finally, while our data shows an important role for ER-Ca^2+^ depletion and NFAT activation in G1-T-cells, TCM expansion was not completely inhibited with calcineurin inhibition and may suggest a role for other signaling pathways downstream of APOL1(50), which need to be investigated in future work. For instance, VEGF levels were altered in supernatant analyses, and VEGF-VEGFR signaling pathway was also enriched in G1-CD8+T-cell subsets. The specific role of this signaling pathway, among others, in APOL1-RV-CD8+T-cells needs to be explored in future work.

In conclusion, we demonstrate for the first time that APOL1-RVs in T-cells induce CD8+T-cell activation by promoting ER-Ca^2+^ leak, impinging on SOCE and canonical TCR signaling. Our work demonstrates a causal role for these variants in promoting AR risk after kidney transplantation, providing opportunities for precision therapeutics directed at carriers of APOL1-exonic variants.

## MATERIALS AND METHODS

**(See Supplemental methods for details)**

### Sex as a biological variable

Animals of both sexes were included, and similar findings were reported in this study.

### BAC transgenic mouse model

Reference APOL1-BAC-Tg G0 construct was initially cloned, and G1, G2-BACs were generated by site-directed mutagenesis. BAC-Tg mice were generated by pronuclear injection at the Yale O’Brien Center Transgenic Core Facility following a method described by Wang et al. (51). Founders with 2 copies of APOL1-BACs were genotyped and backcrossed to the C57B6/J background. Genotyping: Mouse APOL1 copy number was determined using cellular DNA via ΔΔCt method and mouse CT values were compared against human homozygotes. RPS18 was used as an endogenous control for mouse samples while B2M was used for human samples.

### Animal studies

### Ex vivo mouse studies

Mouse spleen and mesenteric lymph nodes are harvested and processed. Whole and naive CD4+ and CD8+ T-cells were isolated from the spleens using EasySep mouse T-cell isolation kits, stained with Cell trace Violet (CTV), and proliferated with anti-CD3/CD28. The effects of an APOL1 inhibitor (MZ-302) on T-cell proliferation and cytokine production, and the effects of BAPTA-AM (at 0.3 and 3.0 uM), YM-58483 (at 50 nM), or Tacrolimus (at 5 and 10 ng/ml) were examined on T-cell proliferation in respective experiments. Intracellular cytokine production was assayed using the BD Cytofix/Cytoperm Plus kit with BD GolgiPlug (BD Biosciences, 555028) after PMA/Ionomycin and BD GolgiPlug.

#### Flow cytometric analysis

The harvested cells were first stained with Zombi NIR and the surface markers and followed by fixation and permeabilization using the eBioscience Foxp3/Transcription Factor Staining Buffer set (Invitrogen, 00-5523-00) to stain the intracellular markers including the transcription factors. The details of tissue processing, T-cell isolation and proliferation, staining protocols, and a list of antibodies and fluorochrome colors used in this study are provided in the supplementary methods. All FACS samples were run on a BD Symphony A5 Cell Analyzer.

#### Thapsigargin-mediated cytosolic calcium assay

CD8+ T-cells were stimulated on coated 96-well plates for 48 hours. To estimate the total ER Ca^2+^-content, T-cells were incubated with Fluo-4 AM and washed with Ca^2+^- free phenol red-free RPMI media containing 2% FBS and 1 mM EGTA (Ca^2+^ chelator). Cells were then stimulated with 10 uM Thapsigargin (Invitrogen# T7459).

#### Anti-CD3-mediated cytosolic calcium assay

For intracellular Ca^2+^ chelation, BAPTA-AM was loaded alongside Fluo-4 AM. Fluorescence intensity (20X) on a Bruker Opterra II Swept-field confocal microscope. Baseline fluorescence was recorded for 20 seconds and then anti-CD3 was added to the cells (5 ug/mL concentration). Changes in Fluo-4 fluorescence were recorded for an additional 3.5 minutes and quantified using ImageJ. Increases in [Ca^2+^]_cyt_ were expressed as percent increase in Fluo-4 fluorescence intensity and normalized by baseline fluorescence.

### In vivo mouse studies

Mice were injected once retro-orbitally with 200 micrograms of poly(I:C) dissolved in PBS (52). Sera were collected at days 1 and 7 post-injection. Spleens were harvested on day 7 post-injection for analysis.

### Murine cytokine assay

Sera cytokines levels from *in vivo* poly(I:C)-treated mice and in the supernatants of *ex vivo* CD8+T-cell proliferation experiments were investigated by Eve Technologies (Calgary, Alberta, Canada) Mouse Cytokine/Chemokine Discovery Assay.

### In vitro studies

APOL1 plasmid constructs kindly provided by Dr. Waldemar Popik (Meharry Medical College) were transiently overexpressed in HEK293T (53). HEK293T cells were also co-transfected with either the VA- or VC, G0-, G1-, or G2 constructs and lentiviral packaging plasmids to generate mammalian VSV-pseudotyped lentiviral expression constructs as previously described (54).

#### Cytotoxicity assays

PI/Annexin V staining was performed using the Pacific Blue Annexin V Apoptosis Detection Kit with PI. WST-1 staining was performed using the Cell Proliferation Assay Kit (Millipore, 2210).

#### Immunoblotting

HEK293T cells and mouse cells were processed to quantify the expression of APOL1. The antibodies and quantification details are provided in Supplemental methods.

#### qPCR

Primer sets were designed for all assayed genes via Primer-BLAST (NCBI) (Primers are listed in Supplemental Table 8). Gene expression was assayed by RT-PCR (Applied Biosystems 7500) via the ΔΔCt method and using GAPDH as endogenous control.

### Transplantation experiments

#### Bone marrow chimera

Bone marrow cells were prepared from G0 and G1 animals (10-12 weeks old), by mechanically flushing and RBC lysis. Recipient animals received whole body irradiation (900 cGy) in a gamma irradiator followed by injection of 5 x10e6 bone marrow cells intravenously. Donor chimerism was confirmed using congenic markers.

#### Heterotopic heart transplants

Heterotopic heart transplant experiments were performed as reported before (55, 56). Recipients were either treated with CTLA-4 Ig (Abatacept, 10 mg/kg i.p. ∼ 250 mg on day 2). A mixed lymphocyte reaction was performed using irradiated donor splenocytes. Recipient cells were stained with CTV, then donor and recipient cells were co-cultured in 24-well plates for 96 hours at 1×10^6^ and 1×10^5^ cells/well. Positive controls were mouse T-cell activator beads) at a bead-to-cell ratio of 1:1) and negative controls (without stimulation). The transplantation experiments are described more in the supplemental method.

### Transcriptome analyses

#### Bulk RNA sequencing

Poly-A library preparation was done at Yale Center for Genome Analyses (YCGA). The quality control was conducted with FastQC v0.11.8, and the adapter sequences trimmed and aligned to the mouse genome (mm39). The differentially expressed gene (DEG) analysis was conducted by limma-voom (V3.46.0). Genes with nominal p value less than 0.01 were identified as DEGs and enrichment analysis was conducted with “enrichR” (V2.1) R package.

#### Single-cell RNA sequencing analysis

Single-cell RNA-seq experiments (library preparation, hashtagging (Biolegend), cDNA synthesis) were performed by the Center for Cellular Profiling at Brigham and Women’s Hospital (10X Genomics protocol). Cells were identified as sample by the highest expressed hashtag sequence. Unsupervised clustering was conducted with the first 30 PCs with a resolution parameter as 0.8. The annotation was based on markers in previous publications (Figure 6D-E). The DEG analysis was conducted with Seurat (V4.1.1). DEG analysis for each cell type was conducted using the Wilcoxon Rank Sum test. Sensitivity analysis was conducted by removing the Hashtag5 sample. Down-sampling DEG analysis was conducted by randomly choosing the same number of G0 cells from the G1 cell population (equally from each G1 sample) for each cell type. Down-sampling was conducted 100 times and DEG consistently seen (sample direction of fold change) in 50 of the 100 analyses were considered robust DEGs. Enrichment analysis was then conducted with “enrichR” (V2.1) R package. Additional information is provided in the supplemental method.

### Human cohorts

APOL1-RV was defined as G1/G1, G1/G2, G2/G2, G1/G0 or G2/G0 genotypes.

#### CHARM study

The details of this longitudinal study on epidemiology, and transmission of SARS-CoV-2 infection of the United States Marine recruits at Marine Corps Recruit Depot, Parris Island, in South Carolina are reported elsewhere (57). The O-link data was downloadable at the publication, while the peripheral blood RNA-seq data was downloadable at GEO [GSE198449].

#### COVID OLINK protein data analysis in humans

The OLINK protein expression profile and demographics of CHARM cohort were freely downloadable (58). The APOL1 genotype was evaluated by aligning the RNA sequencing reads to the APOL1 gene.

#### CTOT-19 study

Related information is described in the supplemental method and the details of this trial including inclusion/exclusion criteria, and principal findings have been reported elsewhere (40).(PI Dr Heeger, Mount Sinai).

#### Genotyping of CTOT-19 recipients

Recipient DNA was genotyped (Illumina GSA3), and imputed (https://imputation-server.sph.umich.edu). APOL1 risk alleles were directly genotyped without imputation. The allele that does not carry any of these variants is identified as G0 (detailed in Supplemental methods). Of 225 total recipients in CTOT-19, 15 recipient genotypes could not be resolved (IRB #2000030257).

#### Flow cytometry CTOT-19

Surface staining was performed on freshly isolated PBMCs using the CTOT protocols as previously published (59). Out of the 56 patients with available FACS data from visit 0, 1 G0/G0 NAR, 1 G0/G0 AR, 2 APOL1-RV NAR, and 1 APOL1-RV AR sample were excluded due to poor staining.

### Statistical analyses

Unpaired t-tests or Mann–Whitney tests, One-way ANOVA (with Tukeys post hoc test) were applied (two-tailed P<0.05). For CTOT-19, descriptive statistics were compared using the chi-squared test and Fisher’s exact test or T-tests. Kaplan–Meier survival curves were calculated with AR as outcome. Bivariate cox proportional hazard models were developed using APOL1-genotype status and covariates that were significantly different between APOL1-RV- and G0/G0 recipients (GraphPad Prism, CA, USA).

### Data Availability

The transcriptomic data included in this work are available on the Gene expression Omnibus repository under GSE289030 (Bulk RNA) and GSE289032(Single-cell RNA). The programming code used is located at https://github.com/ZephyrSun03/APOL1_seq. Detailed CTOT-19 and CHARM clinical data are under the respective publications (40, 60). The CTOT-19 recipient genotyping data is available upon request to the corresponding author. MZ-302 availability is restricted and was obtained through a material transfer agreement between Yale University and Maze Therapeutics, Inc.

## Supplementary Materials included

Supplemental Methods

Supplemental Figures 1-10

Supplemental Tables 6-8

References 01 to 19

Data file (includes supplemental tables S1-S5)

## Acknowledgments

We thank Prof Anita Chong (University of Chicago) for critical discussions regarding the manuscript, and Anushree Vashist for her technical help. We acknowledge Dr Deepika Kumar from Yale pathology for photomicrographs. We thank Dr Terry Satterfield, Dr Julie Ullman, Dr Victoria Assimon, and Maze Therapeutics, Inc. for their collaboration and providing the MZ-302.

We thank Yale Flow Cytometry for their assistance with panel design and sample analysis. The Yale flow cytometry Core is supported in part by an NCI Cancer Center Support Grant # NIH P30 CA016359. The BD Symphony was funded by shared instrument grant # NIH S10 OD026996.

## Author contributions

Conceptualization: MCM

Data Curation: JP, ET, ZS, MCM

Formal Analysis: JP, ET, ZS, MCM

Funding Acquisition: PSH, JA, SI, MCM

Investigation: JP, ET, IC, AR, MG, JC, SC, HM, RI, BF, IG, JA, NM

Methodology: SN, XT, SS, SI, MCM

Project Administration: JP, ET, HS, MCM

Resources: JP, ET, IC, AR, SN, XT, PC, AK, GB, WP, JC, JA, NM, SI, PH, MCM

Supervision: IC, MG, BK, WS, JSP, ZZ, JC, JA, NM, SI, PH, MCM

Validation: JP, ET, ZS, JA, NM, SI, MCM

Visualization: JP, ET, MCM

Writing – original draft: JP, ET, MCM

Writing – review and editing: All co-authors

