## Supplementary Material for "A T-cell intrinsic Role for *APOL1* Risk Alleles in Allograft Rejection"

### List of Supplementary materials

**Materials and Methods**

**Supplemental Figures 1-10**

**Supplemental Tables 6-8 (Supplemental tables 1-5 in Supplemental Data Excel)**

**References**

**Supplemental Methods:**

**Study Design:** The objective of this study was to mechanistically evaluate the role of APOL1 risk variants in T-cells to provide a causal explanation for their association with kidney transplant rejection. Mechanistic studies on T-cells were performed using a BAC transgenic mouse model that allows for physiologic expression of APOL1 in mice under the human promoter. Mice were kept at the Yale animal care facility and animal studies were per IACUC protocol (# 2023-20495). T-cell response to allo-antigen stimulation was evaluated both in vivo and in vitro. Calcium assays were performed on isolated murine CD8+T-cells in vitro. Surface phenotypes as well as cytokine and transcription factor expression were evaluated by flow cytometry and multiplex assays. Transplant studies were conducted using heterotropic mismatched heart transplants in C57Bl/6J APOL1 bone marrow chimeras. Chimeras were generated and housed in the Brigham and Women's Hospital animal care facility per ICAUC protocol #2019N000153. In vitro studies using 293T cells were conducted using DOX-inducible APOL1 lentiviral constructs. Human data was analyzed from both the CHARM and CTOT-19 studies (study designs and data collection described in supplemental methods). Demographic information for these studies can be found in Tables 1 and 2, respectively.

**Generation and genotyping of BAC transgenic mouse model**

APOL1-BAC transgenic mice (BAC-Tg) were generated by pronuclear injection (and random genomic integration) using a 104 kb BAC bacterial artificial chromosome (BAC) that includes the APOL1 gene with partial fragments of the APOL2 and MYH9 genes (G0, G1, G2). The BACs were generated using homologous recombination in E. coli with a positive/negative selection marker, as described elsewhere (1). The genotype of the original BAC was G0 ENSEMBLE ID; G1 and G2 BACs were subsequently generated via site-directed mutagenesis (S342G and I384M for G1, deletion of aa 388-389 for G2, Yale O'Brien Center Transgenic Core Facility). Following pronuclear injection of each BAC into fertilized C57Bl/6 oocytes, and generation of founders, founders with very low and very high APOL1 BAC copy numbers (in comparison to human DNA which has 2 copies of APOL1 gene) were deselected. We focused on copy numbers since expression of BAC genes is copy number dependent, but relatively independent of integration site. These founder lines were maintained by Dr Shuta Ishibe, and were obtained from the Ishibe lab for splenocyte studies. Due to varying coat colors among offspring, G1 and G0 lines were progressively backcrossed into C57B6/J lines [Jackson lab #000664]. Mice with

~2 copy numbers (comparing with human genome using genomic qPCR [Supplemental Figure 1A]) were selected at each step both for experimentation and backcrossing. Genomewide C57B6/J genotype after backcross was confirmed using Mini-MUGA SNP-array (Transnetix, inc).

**DNA qPCR**

For mouse copy number determination, tissue samples were homogenized and lysed, then cellular DNA was isolated using the QIAamp DNA Micro Kit (Qiagen, 56304). Gene expression was assayed by RT-PCR (BioRad CFX96). Amplification curves were analyzed via the ΔΔCt method. Mouse sample CT values were compared against human homozygotes. RPS18 was used as an endogenous control for mouse samples while B2M was used for human samples.

**Animal husbandry and breeding**

Animals were housed according to protocol (IACUC protocol # 2023-20495). All animal lines were housed in the same room in separate cages. Mice were used for experimentation at around 8-10 weeks of age. All practices involving animals were performed in compliance with the Institutional Animal Care and Use Committee (IACUC) at Yale University.

**<u>Ex vivo mouse studies:</u>**

**Splenocyte isolation**

Mice were euthanized according to IACUC protocol. Following euthanasia, whole spleens were harvested and processed by passing them through a 70uM cell strainer with a 10 mL syringe. Splenocytes were de-hemoglobinized with ACK lysis buffer (Gibco, A10492-01) for five minutes at room temperature at a concentration of 5 mL per spleen.

**Ex vivo Mouse T-cell culture**

Whole and naive mouse CD4+ and CD8+ T-cells were isolated using Stem Cell EasySep mouse T cell isolation kits (19853,19858 and 19852, 19765 respectively). T-cells were then stained with Cell trace Violet (CTV) (Invitrogen, C34557) and proliferated in a 96-well plate (Falcon, 353077) with 200 uL/well of complete RPMI media (+ beta mercaptoethanol, 0.1%) with anti-CD3/CD28 coated beads (Miltenyi Biotec, 130-093-627) at a bead to cell ratio of 1:2. At 72 hours, the supernatants were collected, and cells were stained with TCR-beta, CD4, CD8, CD44, CD69, CCR7, KLRG1 (2F1/KLRG1, FITC, 1:200), and a Zombie NIR viability stain. Inhibitor studies included a novel APOL1-inhibitor (10 uM, Maze Therapeutics, inc.), BAPTA-AM (B6769, Invitrogen, 0.3 and 3.0 uM), CRAC channel inhibitor (YM-58483, Abcam, 50 and 300 nM). YM-58483 was replenished during 72-hour proliferation.

**Intracellular Markers and Transcription Factor staining**

Fixation/permeabilization was performed using the eBioscience Foxp3/Transcription Factor Staining Buffer set (Invitrogen, 00-5523-00). Following surface marker and viability staining (TCRbeta, CD4, CD8 (clone 53-6.7, BV650, 1:200, 100741), CD44, CD69, KLRG1 (2F1/KLRG1, FITC, 1:200, 11-5893-82), CCR7, CD183, Zombie NIR), cells were fixed and permeabilized at room temperature for one hour using the kit. Cells were stained at room temperature for 30 minutes with Foxp3 (clone R16-715, PerCP-Cy5.5, 1:100, 563902), RORgT (clone Q31-378, BV480, 1:100, 567176), EOMES (clone X4-83, BUV737, 1:100, 567170), Tbet (clone 4B10, PE, 1:100, 561265), GATA3 (clone L50-823, BV711, 1:100, 565449), Blimp1 (clone 5E7, APC, 1:100, 2311257), and TCF-7/TCF-1 (clone S33-966, BV421, 1:100, 566692), Perforin (clone eBioOMAK-D, APC, 1:100, 17-9392-80)

**Intracellular cytokine staining**

Intracellular cytokine staining was performed using the BD Cytofix/Cytoperm Plus kit with BD GolgiPlug (BD Biosciences, 555028). Following surface marker and viability staining, cells were stimulated with PMA/Ionomycin and incubated with BD GolgiPlug for 5 hours, then fixed and permeabilized for 20 minutes at 4 degrees C using the kit. Cells were stained at room temperature for 30 minutes with TNFa (clone MP6-XT22, BV480, 1:100, 414-7321-82), IL-1beta (clone NJTEN3, FITC, 1:100, 11-7114-82), IL-2 (clone JES6-5H4, BUV737, 1:100, 367-7021-82), and IFNG (clone XMG1.2, BV605, 1:100, 505839).

**Flow Cytometry**

For surface marker analysis of adaptive immune cell subsets, samples were stained with TCRbeta (clone H57-597, APC, 1:200, 553174), CD19 (clone 1D3, PE-Cy7, 1:200, 25-0193-82), CD4 (clone RM4-5, AF700, 1:200,

100536), CD8 (clone 53-6.7, BV711, 1:200, 100759), CD44 (clone IM7, PE-Dazzle 594, 1:200, 103056), CD69 (clone H1.2F3, BUV395, 1:100, 569367), CD23 (clone B3B4, PE, 1:200, 553139), and IgD (clone 11-26c.2a, BV605, 1:500, 405727). To evaluate innate immune cell subsets, samples were stained with TCRbeta, CD19, CD49b (clone HMa2, BV421, 1:200, 740030), NK1.1 (clone PK136, BV711, 1:200, 740663), CD14 (clone rmC5-3, FITC, 1:200, 123307), and DCIR2 (clone 33D1, PE, 1:200, 557578). To evaluate T-cell subsets, samples were stained with TCRbeta, CD4, CD8, CD44, CD69, CCR7 (clone 4B12, PE-Cy7, 1:50, 120124), KLRG1 (clone 2F1/KLRG1, BV421, 1:200, 138414), CD183 (clone CXCR3-173, BUV661, 1:100, 741681). All samples were stained with a Zombie NIR viability stain (APC-Cy7, 1:1000, 423106). CCR7 was stained at room temperature for 30 minutes, all other markers were stained at 4 degrees C. All FACS samples were run on a BD Symphony A5 Cell Analyzer.

**Realtime measurements of anti-CD3-mediated cytosolic calcium $[Ca^{2+}]_{cyt}$ in CD8+ T cells:**

Isolated CD8+ T-cells (20,000 cells/well) were cultured for 48 hours on anti-CD3/CD28 (2.5 ug/mL) coated 96-well glass bottom plates (Cellvis# P96-1.5H-N) with complete RPMI media (+beta mercaptoethanol, 0.1%). The culture media was, then removed and replaced with phenol red-free RPMI media (Gibco# 11835030) containing 5 uM of Fluo-4 AM (Invitrogen #F14201), supplemented with 2% FBS and 2 mM $CaCl_2$ (Sigma# C-34006), and cells were incubated for 30 minutes at 37° C and 5% $CO_2$. For intracellular $Ca^{2+}$ chelation, BAPTA/AM (Invitrogen# B6769) was loaded alongside Fluo-4 AM. The real-time fluorescence intensity was directly monitored at 20X magnification on the stage of a Bruker Opterra II Swept-field confocal microscope (Billerica, MA) equipped to maintain 37° C and 5% $CO_2$ at 1 Hz. The baseline fluorescence was recorded for 20 seconds and then anti-CD3 was added to the cells at a final concentration of 5 ug/mL prepared in the same media. Changes in Fluo-4 fluorescence were recorded for an additional 3.5 minutes and quantified using ImageJ. Increases in $[Ca^{2+}]_{cyt}$ were expressed as percent increase in Fluo-4 fluorescence intensity and normalized by baseline fluorescence.

**Realtime measurements of Thapsigargin-mediated cytosolic calcium $[Ca^{2+}]_{cyt}$ in CD8+ T cells:**

To estimate the total ER $Ca^{2+}$-content, T-cells were incubated with Fluo-4 AM and washed with $Ca^{2+}$-free phenol red-free RPMI media containing 2% FBS and 1 mM EGTA (Millipore cat # 324626). Cells were then stimulated with 10 uM Thapsigargin (Invitrogen# T7459) prepared in $Ca^{2+}$-free media (by chelation with EGTA (ethylene glycol bis(2-aminoethyl ether)-N,N,N)).

**In vitro studies:**

**HEK 293T cell culture and transient overexpression studies**

HEK 293T cells were cultured in DMEM supplemented with 10% FBS, 1% penicillin-streptomycin and 0.1% fungin in a CO2 incubator (Thermo, 13-998-253) maintained at 37$^{o}$ C and 5% CO2. Cells were incubated in a media depth of ~0.172 cm and cultured at a cell density of ~140,000 cells/cm$^2$. APOL1 plasmid constructs were kindly provided by Dr. Waldemar Popik (Meharry Medical College, Nashville, USA) and have been previously published (2). Transient transfection of APOL1 constructs (6 ug each) was carried out in HEK293T cells at 70% confluence using Turbofect transfection reagent (Thermo, R0531) overnight. Following transfection, APOL1 overexpression was induced with doxycycline (100 ng/mL). Overexpression was confirmed via western blot and qPCR. RNA for qPCR was isolated using the RNeasy Mini Kit (Qiagen, 74106) with DNAase cleanup (Qiagen, 79254) to eliminate plasmid contamination.

**Lentiviral vector generation**

HEK293T cells were used as producer cells for lentiviruses. The cells were transfected with 4 µg each of either the VA- or VC, G0-, G1-, or G2 constructs (obtained from Dr Waldemar Popik(2)). Lentiviral packaging plasmids (pLenti CMV/TO Puro DEST), pPACK, and pVSV using TurboFect transfection reagent (Thermo Scientific #R0531) were co-transfected to generate mammalian VSV-pseudotyped lentiviral expression constructs as previously described (3) The media containing lentiviral constructs were collected, filtered and concentrated using Lenti-X Concentrator (#631231; Takara) following the manufacturer’s protocol. These lentiviral constructs were used to infect HEK293T-cells and the stably overexpressing cells were selected by puromycin (#ant-pr-1, InvivoGen) treatment. APOL1 expression was then induced by Doxycycline treatment at 100 ng/mL concentration.

**Cytotoxicity Assays**

PI/Annexin V staining was performed using the Pacific Blue Annexin V Apoptosis Detection Kit with PI (BioLegend, B415797) according to the kit's instructions. WST-1 staining was performed using the Cell Proliferation Assay Kit (Millipore, 2210). Cells were cultured at a density of 0.1-5x$10^4$ cells/well in a 96-well microplate for 24 hours. WST-1/ECS solution was diluted 10X and added at 100 uL/well, then cells were incubated for 30 minutes in a 37° C, 5% CO2 incubator. Cells were then placed for one minute on a shaker and absorbance was read on a microplate reader at 450/630 nm.

**Western Blotting**

HEK293T cells and mouse cells were lysed using RIPA buffer added with Halt Protease and Phosphatase Inhibitor (Thermo Scientific, 78440) as mentioned elsewhere. Cells were resuspended in buffer and incubated on a rotator at 4°C for 1 hour. The lysate was then centrifuged at 12000xg at 4°C and the supernatant was collected. The protein concentrations of the supernatant lysates were determined by BCA assay (#23225, Pierce-Thermofisher). Following BCA quantification, loading buffer was added and the protein lysates were denatured at 95°C for 5 minutes. The proteins were separated on 4-15% gels (Bio-Rad) by SDS-PAGE and transferred to an ImmunoBlot PVDF membrane (BioRad, 1620177) followed by blocking with 5% skim milk. The primary antibodies used were anti-APOL1 (Sigma, HPA018385), β-Actin (Sigma, A5441), anti-APOL1 (Proteintech, 66124-1-Ig), anti-HSP-90 (Cell Signaling, 4877). The secondary antibodies used for detection included horseradish peroxidase (HRP)-conjugated anti-rabbit and anti-mouse antibodies, with a 1:10000 dilution. Image Studio Lite Ver5.2 was used for western blot image acquisition. Densitometry was performed on images of Western blots using ImageJ software.

**mRNA reverse transcription qPCR**

Primer sets were designed for all assayed genes via Primer-BLAST (NCBI). Gene expression was assayed by RT-PCR (Applied Biosystems 7500). Amplification curves were analyzed using the automated 7500 software platform via the ΔΔCt method. GAPDH was used as an endogenous control.

**<u>Transplantation experiments:</u>**

**Bone marrow chimera:**

Bone marrow cells were prepared from femur and tibia bones obtained from G0 and G1 animals (age 10-12 weeks old). Briefly, bone marrow cells were mechanically flashed out of bones using 27-gauge insulin syringes, then red blood cells were lysed with ACK lysis buffer (Gibco) for 2 minutes on ice. One donor animal was sufficient to perform transplantation into 4-5 bone B6/45.1 recipient mice (all males). Recipient animals received whole body irradiation (900 cGy) in a gamma irradiator (Gammacell 40 Exactor, Best Theratronics Ltd.), on the day of transplantation, 6 hrs prior, followed by injection of bone marrow cells (5 x10e6 cells per animals) intravenously. Donor chimerism was confirmed by flow cytometry at 4-6 weeks after bone marrow transfer, using peripheral blood samples from recipient animals, stained with CD45.1 (clone A20, Biolegend) and CD45.2 (clone 104, Biolegend) congenic markers.

**Heterotopic heart transplants**

Heterotopic heart transplant experiments were performed as reported before (4, 5). Donor and recipient animals were anesthetized with isoflurane, and donor hearts were removed and transplanted as follows. The recipient's infra-renal abdominal aorta and inferior vena cava were isolated and cross-clamped, and the donor aorta and pulmonary artery were joined end-to-side to the recipient aorta and vena cava, respectively. The abdomen was closed after confirming the beating donor heart. Recipients were treated with either CTLA-4 Ig (Abatacept, 10 mg/kg i.p. ~ 250 mg on day 2 post-transplant). Graft survival was monitored daily through palpation by a researcher blinded to the experiment. Rejection was defined as the day on which a palpable heartbeat was no longer detectable.

**Immunologic assessment**

Mixed lymphocyte reaction was performed using irradiated donor splenocytes. Recipient cells were stained with CTV, then donor and recipient cells were co-cultured in 24-well Corning plates for 96 hours (37° C, 5% CO2 incubator) at $1x10^6$ and $1x10^5$ cells/well, respectively. Following incubation, FACS analysis was performed for surface activation markers and CTV dilution. Positive control was incubation with mouse T-cell activator beads

(Thermo, 11453D) at a bead to cell ratio of 1:1. Negative control was performed with absence of any stimulation in otherwise identical conditions.

**Transcriptome analyses:**

**Bulk RNA sequencing analysis**

RNA quality was checked using RIN numbers and nanodrop. Library preparation of quality-affirmed RNA from each sample was done using poly-A selection at Yale Center for Genome Analyses (YCGA). The quality control of the raw sequencing data was conducted with FastQC v0.11.8 and the adapter sequences were trimmed by cutadapt (V4.1)(6) with parameters as "-a AGATCGGAAGAGCACACGTCTGAACTCCAGTCA -A AGATCGGAAGAGCGTCGTGTAGGGAAAGAGTGT". The cleaned sequences were aligned to the mouse genome (Ensembl mm39) using STAR v2.7.5.b(7) with default parameters. The expression was quantified as the read counts within each gene region using htseq-count with parameters, "-m union --nonunique none". The differentially expressed gene (DEG) analysis was conducted by limma-voom (V3.46.0) (8) with read count matrix, adjusting for experiment batches. Genes with nominal p value less than 0.01 were identified as DEGs and the enrichment analysis was conducted with "enrichR" (V2.1) R package(9) with Gene Ontology (GO) (2021), WikiPathways (2021) and KEGG (2021) database. (6) with parameters as "-a AGATCGGAAGAGCACACGTCTGAACTCCAGTCA -A AGATCGGAAGAGCGTCGTGTAGGGAAAGAGTGT". The cleaned sequences were aligned to the mouse genome (Ensembl mm39) using STAR v2.7.5.b(7) with default parameters. The expression was quantified as the read counts within each gene region using htseq-count with parameters, "-m union --nonunique none". The differentially expressed gene (DEG) analysis was conducted by limma-voom (V3.46.0) (8) with read count matrix, adjusting for experiment batches. Genes with nominal p value less than 0.01 were identified as DEGs and the enrichment analysis was conducted with "enrichR" (V2.1) R package (9) with Gene Ontology (GO) (2021), WikiPathways (2021) and KEGG (2021) database.

**Single-cell RNA sequencing analysis**

Library preparation was performed per manufacturer's instructions (Single cell 3' v2 protocol, 10x genomics). Single-cell RNA-seq experiments were performed by the Center for Cellular Profiling at Brigham and Women's Hospital. T cells ($CD45^+CD3^+B220^-LiveDead^-$) from the cardiac allografts were isolated using collagenase(5), followed by FACS sorting. Cells from each sample were stained with one of the anti-mouse TotalSeqC cell hashing antibodies (BioLegend). Next, cells from each condition were pooled, and 60,000 cells were resuspended in 0.4% BSA in PBS at a concentration of 1,000 cells per μl and loaded onto a single lane (Chromium chip, 10X Genomics) for encapsulation in lipid droplets, using the Single Cell GEM-X 5' v3 (10X Genomics). cDNA synthesis and gene expression were performed according to the 10X Genomics protocol. The generated gene expression libraries were sequenced to an average of 30,000 reads per cell using the Illumina Novaseq X Plus platform at the Molecular Biology Core Facilities at the Dana-Farber Cancer Institute (DFCI). ScRNA-seq reads were processed with Cell Ranger version 8.0.0, which quantified transcript counts per putative cell. Ambient RNA was quantified and deducted with SoupX (V1.6.2) R package(10). The quality control, clustering and DEG analysis was conducted with Seurat (V4.1.1)(10). Genes expressed in less than 3 cells were removed and cells with less than 200 genes expressed were removed. Potential doublets were identified and removed by DoubletFinder (V2.0.3)(11). Each cell was labeled as the sample with the highest expressed hashtag protein. Unsupervised clustering was conducted with the first 30 PCs with a resolution parameter as 0.8. The annotation was based on markers in previous publications [Fig 6D-E](12, 13). The proliferating T-cell cluster was further divisible into CD8+, CD4+ or Foxp3+T-cells (not shown). DEG analysis for each cell type was conducted by FindMarkers using the Wilcoxon Rank Sum test. Sensitivity analysis was conducted by removing the Hashtag5 sample which had more naïve T cells than others. Down-sampling DEG analysis was conducted by randomly choosing the same number of G0 cells from the G1 cell population (equally from each G1 sample) for each cell type. The down-sampling analysis was conducted 100 times randomly and the genes identified as DEG consistently (sample direction of fold change) in 50 of the 100 analyses were identified as robust DEGs. Enrichment analysis was conducted with "enrichR" (V2.1) R package(9) with Gene Ontology (GO) (2021), Wiki Pathways (2021) and KEGG (2021) database.

**<u>Human cohorts:</u>**

***<u>CHARM study</u>:*** The details of this study are reported elsewhere (14). Briefly, this was a longitudinal study on epidemiology, and transmission of SARS-CoV-2 infection of the United States Marine recruits (all with mild or no symptoms) at Marine Corps Recruit Depot, Parris Island, in South Carolina. Enrollees had weekly nasal SARS-CoV2 PCRs along with blood collections for serology, and cytokine analyses till the end of study. The clinical/demographic data collected for the primary study is located in prior publications and was downloaded without any identifiers (14, 15). The O-link data was downloadable at the publication (15), while the peripheral blood RNA-seq data was downloadable at GEO [GSE198449].

**COVID OLINK Protein Data Analysis in Humans:**

The OLINK protein expression profile and demographic information of the CHARM cohort were generated from their previous publication and are freely downloadable (15). The APOL1 genotype was evaluated by aligning the RNA sequencing reads to APOL1 gene and the genotyping was performed by bcftools mpileup (V1.9). The protein expression difference between groups (G1 vs G0 and (G1 + G2) vs G0) was tested using a linear mixed model with lme4 (V4-1.1) R package along the 0-28 days' time frame. The overall expression of each group was fitted using "loess" with 95% interval.

***CTOT-19 study:*** This was an NIAID funded, phase 2, multicenter, randomized, double-blind, placebo-controlled two-arm study of primary deceased-donor kidney transplant recipients randomized 1:1, to receive intraoperative TNF-blockade with infliximab (single dose of 3 mg/kg) or placebo. This study targeted to enroll 300 recipients but was completed after 225 enrollees. The details of this trial including inclusion/exclusion criteria, basic demographics of the cohort, and principal findings have been reported elsewhere (16). Uniformly, all CTOT-19 patients received Thymoglobulin induction. Surveillance kidney allograft biopsies were performed at 3 and 24 months after transplant. These biopsies were read by a single kidney transplant pathologist (I.W.G. at the central core), and/or reported by each center's local pathologist, and Banff 2013 scores were reported. Indication biopsies were also performed at the discretion of the local investigator. All "for-cause" biopsies were read by the local pathologists for clinical care. All mechanistic flow cytometric studies for T-cells were performed at a central core lab (Dr Heeger, Mount Sinai) at 7 time points through the study. Visit-0 corresponded to the pre-transplant, pre-induction visit for each enrollee (see details in (16)). Genotyping and analyses of CTOT-19 samples were conducted for our study under a separate IRB-approved protocol [#2000030257]. For the purpose of this study, any APOL1-RV was defined as G1/G1, G1/G2, G2/G2, G1/G0 or G2/G0 genotypes.

**Genotyping CTOT-19 kidney transplant recipients**

Recipient DNA was obtained from peripheral blood mononuclear cells (PBMCs) and genotyped with Genome-wide SNP array (Illumina Infinium Global Screening Array; GSA3). We performed genome-wide genotype imputation using the Michigan Imputation Server (https://imputation-server.sph.umich.edu) as described previously (17). We excluded samples with 1) genetically inferred sex not matched with reported sex; 2) missing genotype rate > 0.03; 3) excessive genome-wide heterozygosity, an indication of sample contamination. We excluded SNPs with 1) missing rate > 0.05; 2) minor allele frequency (MAF) < 0.01; 3) Hardy-Weinberg equilibrium (HWE) p- value < 1e-6. The APOL1 risk alleles were directly genotyped from SNP array without imputation. The G1 allele of APOL1 is represented by rs73885319 and rs60910145, two missense SNPs in almost perfect linkage disequilibrium, whereas the G2 allele is represented by a 6 bp microdeletion rs143830837 (or equivalently rs71785313)(18). The allele that does not carry any of these variants is hereafter referred to as G0. Of 225 total recipients in CTOT-19, 15 recipient genotypes could not be resolved either due to failed imputation, failed QC or lack of DNA and were not included.

**Flow cytometry CTOT-19:**

Surface staining was performed on freshly isolated PBMCs from the CTOT-19 study using the CTOT protocols as previously published(19).(19). Samples were stained with the following antibodies: CD8 (clone RPA-T8, BV510, 563256), CD4 (clone RPA-T4, PE-Cy7, 25-0049-42), CD45RA (clone HI100, APC, 550855), CD45RO (clone UCHL1, FITC, 555492) CD27 (clone M-T271, PE, 555441), and CD28 (clone CD28.2, BV421, 56213). Out of the 56 patients with available FACS data from visit 0, 1 G0/G0 NAR, 1 G0/G0 AR, 2 APOL1-RV NAR, and 1 APOL1-RV AR sample were excluded due to poor antibody staining. Three APOL1-RV samples (2 NAR and 1 AR) and one G0/G0 sample (1 NAR) were excluded from analysis on account of poor staining.

**Statistical Analyses**

Statistical analyses were performed in the GraphPad Prism 10 software (Dotmatics, CA, USA). Data are presented as the mean ± SEM. Unpaired t-tests or Mann–Whitney tests were performed for univariate comparisons between two groups. One-way ANOVA (with Tukeys post hoc test) was used while comparing more than 2 groups. Statistical significance was considered with two-tailed P<0.05. For CTOT-19, Descriptive statistics (means and SD) were used to summarize the baseline characteristics of the APOL1-RV- and G0/G0-recipients and were compared using the chi-squared test and Fisher’s exact test. Univariate comparisons of continuous variables were done using unpaired *t* test (Mann–Whitney test for corresponding nonparametric analysis). Kaplan–Meier survival curves were calculated with AR as outcome. Bivariate cox proportional hazard models were developed using APOL1-genotype status and covariates that were significantly different between APOL1-RV- and G0/G0 recipients.

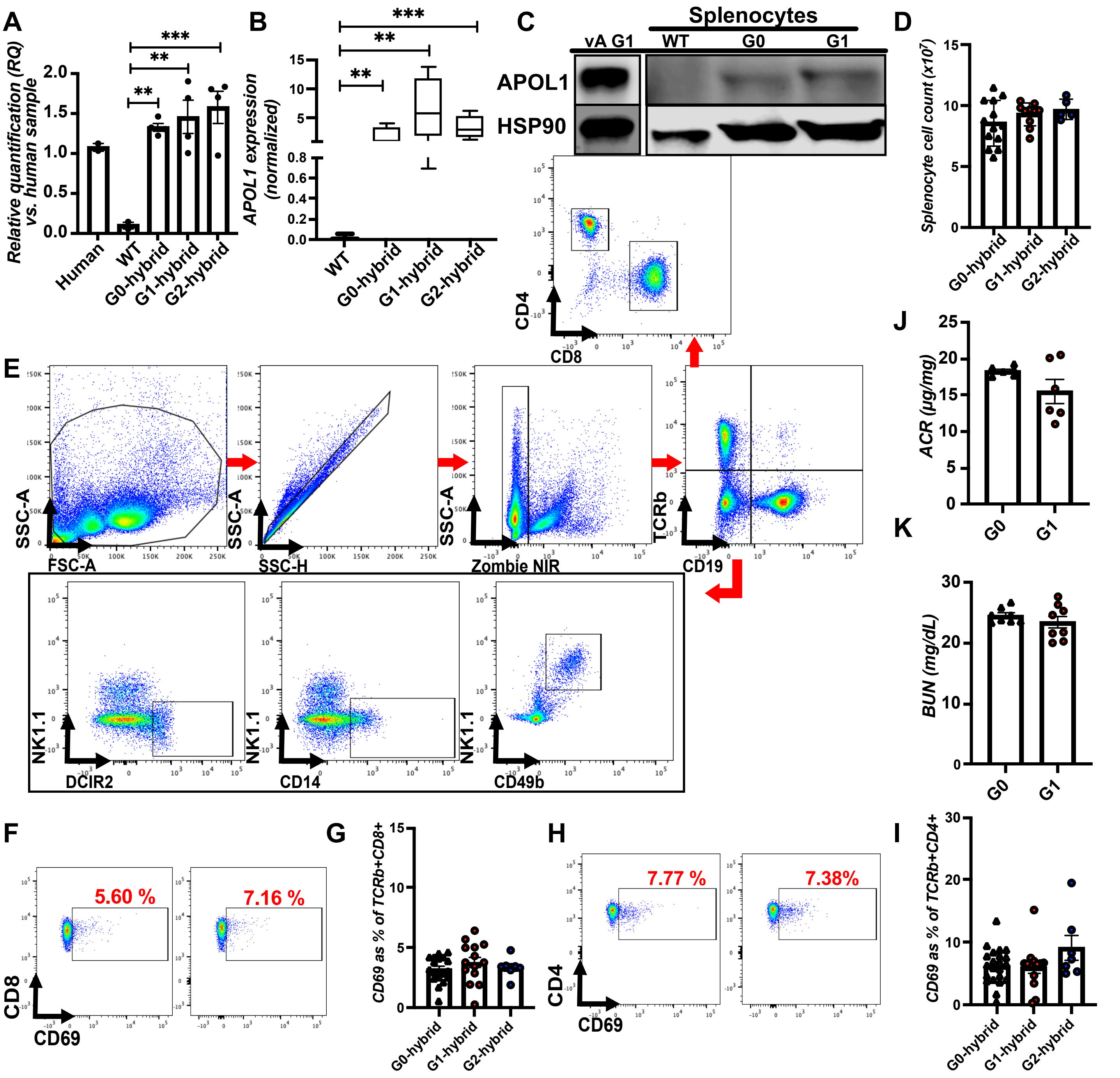

**Supplemental Figure 1: APOL1-BAC transgenic mice development and baseline T-cell phenotypes.** (A) Relative quantification by qPCR of *APOL1-BAC* copy numbers in G0-, G1-, and G2-hybrid mice lines, and the human genome (diploid). (B) Box-whisker plots show qPCR of *APOL1* mRNA from G0-, G1-, and G2-hybrid mouse splenocytes (normalized to *Gapdh*) [n>5 mice each]. (C) Immunoblot of splenocyte lysates from G0- and G1-hybrid mice at baseline probed for APOL1; APOL1 overexpressing HEK293T cells (vA G1) was used as a positive control. (D) Total splenocyte count between G0, G1, and G2 mouse lines at baseline. (E) Flow plots of the gating strategy of the major innate immune cells using markers for dendritic cells (DCIR+), monocyte-macrophages (CD14+), NK cells (NK1.1+), as well as adaptive immune cells including T-cells (TCRb+), CD4+ and CD8+T-cells, and B-cells (CD19+). (F-I) (F) and (H) are representative flow plots while (G) and (I) quantify proportions of CD8+CD69+T-cells and CD4+CD69+T-cells, respectively. Box-Whisker plots show median and range from >5 mice. The bar graphs with mean ± SEM of $n \geq 7$ mice. **$p < 0.01$, ***$p < 0.001$, while circles/triangles represent individual mice.

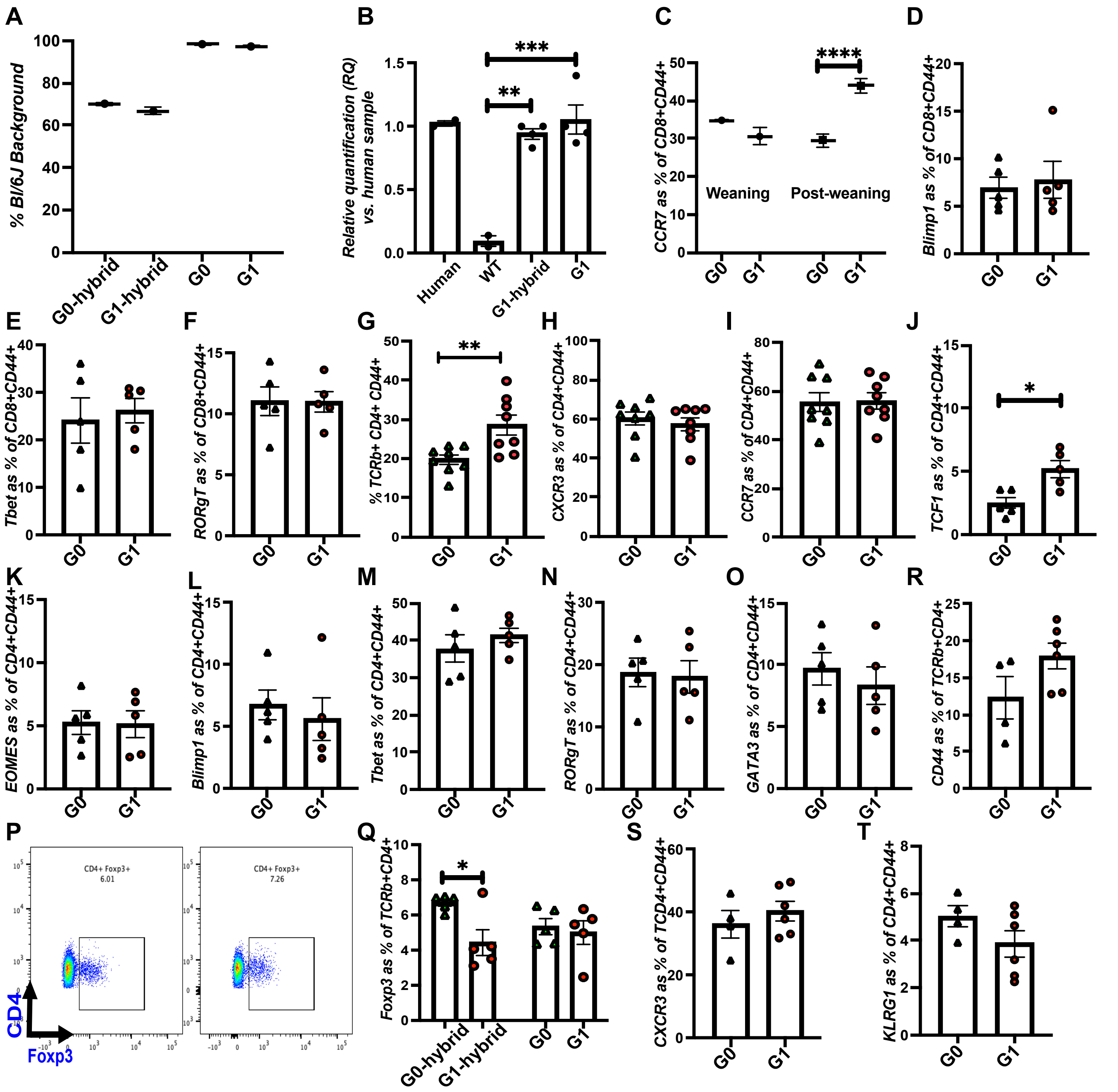

**Supplemental Figure 2: Baseline activation and polarization of CD8+ and CD4+ T-cells in backcrossed variant APOL1-BAC-Tg mice** (A) Box-whisker plots show genetic monitoring by Mini-MUGA SNP-array compares B6-origin SNPs in G1- and G0-hybrid mice, and in G1- and G0- BAC-Tg C57Bl/6 lines (called G1 and G0 after backcrossing) [n>5 mice]. (B) Relative quantification by qPCR of *APOL1* copy numbers in each generation and after backcross (vs human diploid genome). (C) Box-whisker plots show TCM expansion (CD8+CD44+CCR7+) with age, at weaning (4-weeks), and thereafter (until ~20-weeks) [n>5 mice each]. (D-F) FACS analysis of G0 and G1 splenic CD8+CD44+ T-cells after intracellular staining for (D) Blimp1, (E) Tbet, and (F) RORgT. (G) FACS analysis of G0 and G1 splenic CD4+CD44+T-cells proportions. (H-O) Splenic CD4+CD44+T-cells were stained for cell surface polarization markers, (H) CXCR3 and (I) CCR7, as well as intracellular staining for (J) TCF1, (K) EOMES, (L) Blimp1, (M) Tbet, (N) RORgT, and (O) GATA3 (Th2 marker). ((P) Representative flow plot and (Q) proportion of Tregs (CD4+FoxP3+) in G1- and -G0 spleens (pre- and post-backcrossing); (R-T) FACS analysis of mesenteric lymph node for the proportions of CD4+CD44+, CD44+CXCR3+, and CD44+KLRG1+ CD4+T-cells between the G0 and G1 mice. Box-Whisker plots show median and range from >5 mice. The bar graphs with mean ± SEM, n=4-10 mice. *$p < 0.05$, **$p < 0.01$, ***$p < 0.001$, while circles/triangles represent individual mice.

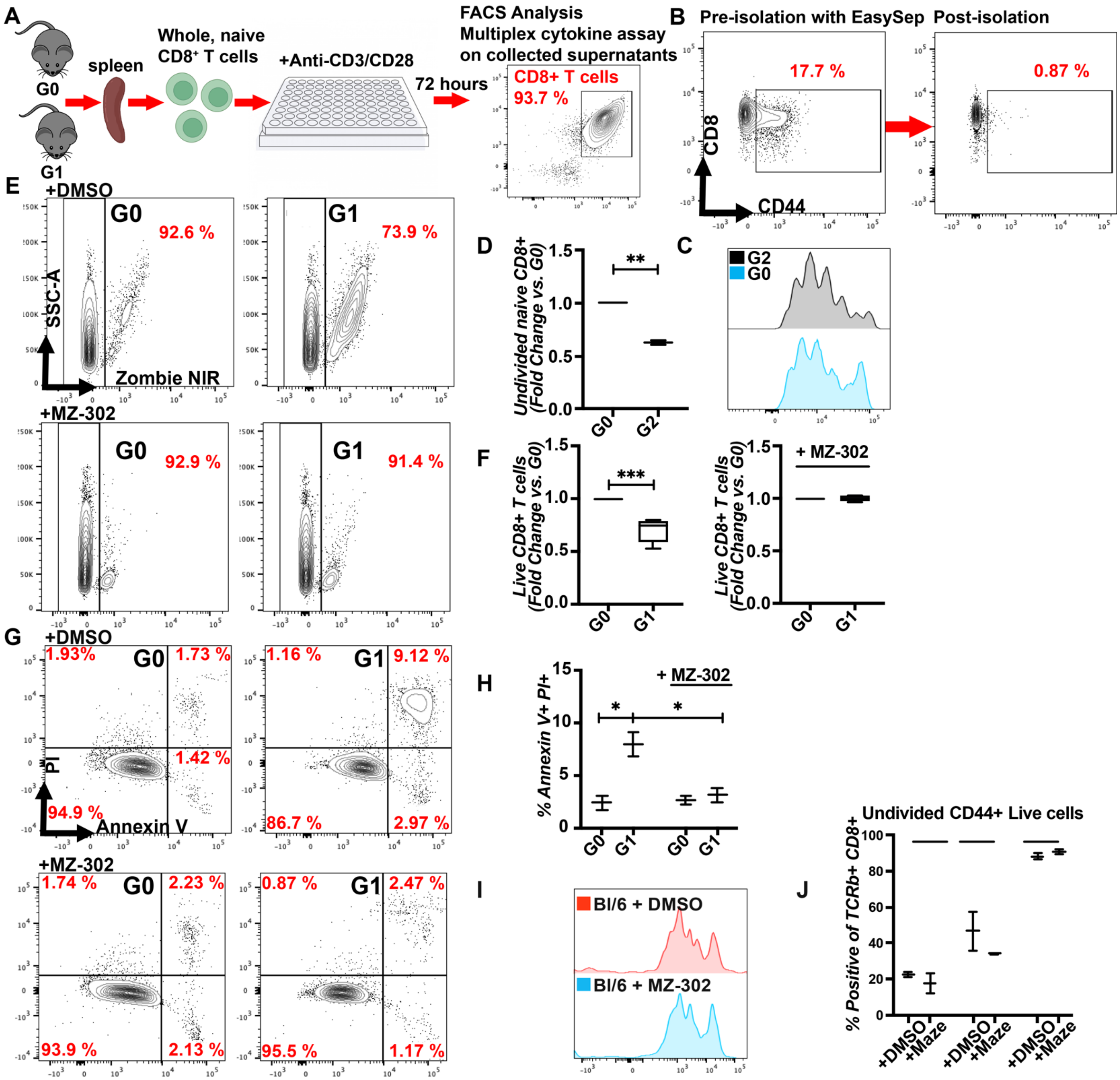

**Supplemental Figure 3: G1-APOL1 CD8+T-cells show increased proliferation and cytokine expression with T-cell receptor stimulation, which is reversed by APOL1 inhibition.** (A) A schematic of splenic CD8+T-cell isolation and *ex vivo* CTV-stained T-cell proliferation with anti-CD3 and anti-CD28 in a 96-well plate for 72 hours (B) FACS analysis shows the high purity of the isolated CD8+T-cell using EasySep kit. (C) Proliferation plot and (D) quantification of undivided native G2-CD8+T-cell (fold change) normalized to G0 (n=5). (E) representative flow plots and (F) quantification of relative T-cell viability with and without MZ-302. (G) Representative flow plots and (H) quantification of apoptotic cell death (Annexin V+ and PI+T-cells) in G1-CD8+T-cells normalized to G0 with and without MZ-302. (I) Representative flow plot and (J) quantification shows absence of effect of MZ-302 on WT Bl/6 T-cell proliferation, activation and viability. Box-Whisker plots show median and range from triplicate wells pooled from 5 mice. *p < 0.05, **p < 0.01, ***p < 0.001.

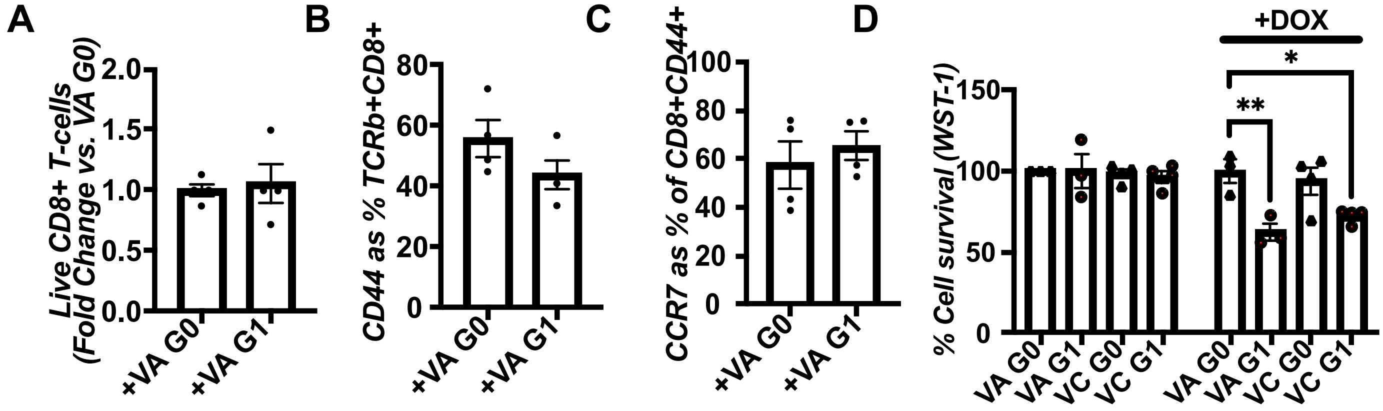

**Supplemental Figure 4: G1-CD8+T-cells show increased proliferation/cytokine expression with TCR stimulation, reversed by APOL1 inhibition.** (A) Live CD8+T-cells and proportions of (B) CD8+CD44+, (C) CD8+CD44+CCR7+T-cells post-supernatant treatment with either VA G0 or G1. (D) Viability of HEK293T cells after inducing VA- or VC- G1 or G0 constructs (n=4 sets). *In vitro* expression of the bar graphs with mean ± SEM represent triplicate wells pooled from 5 mice. *p < 0.05, **p < 0.01.

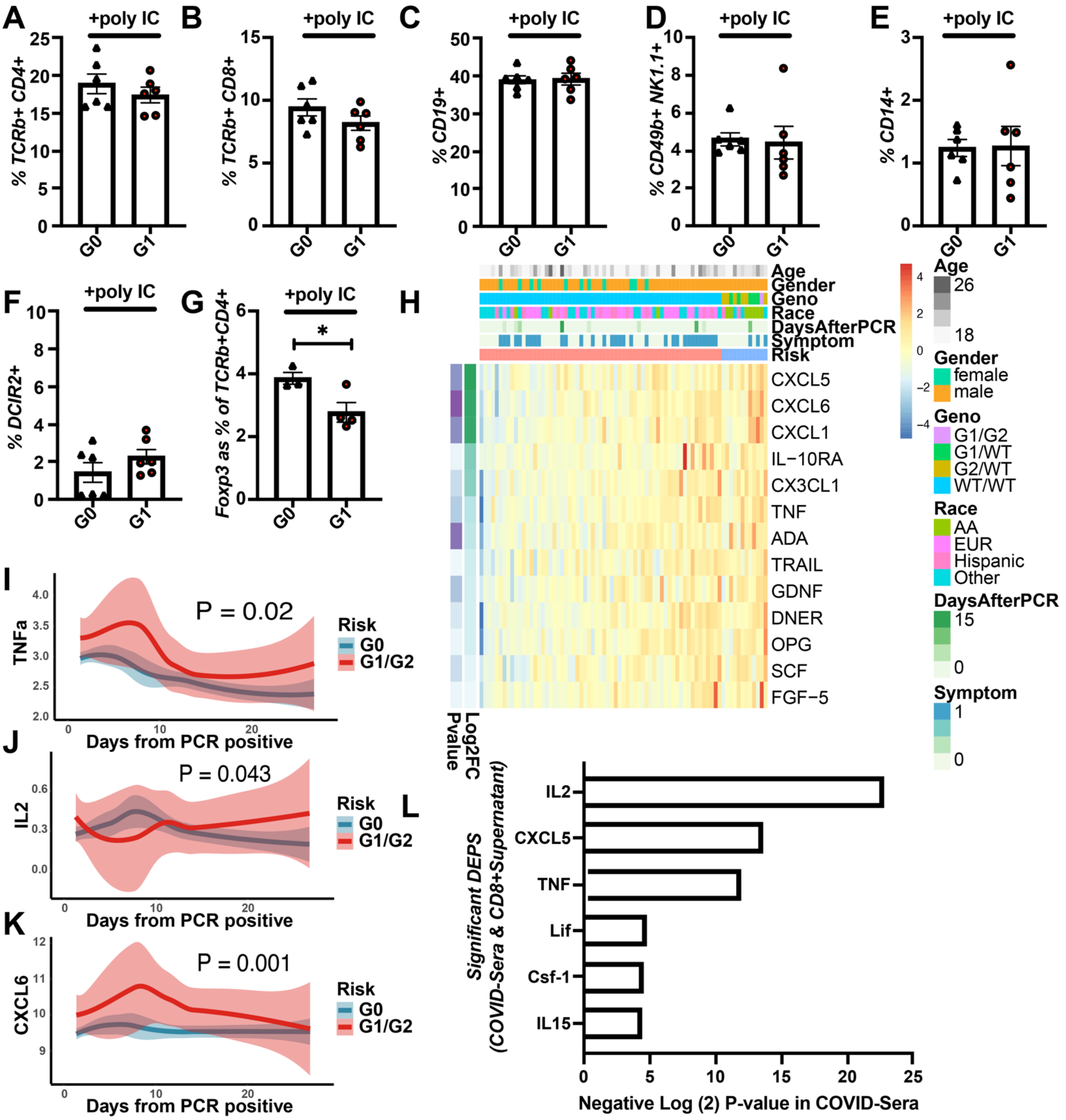

**Supplemental Figure 5: CD8+T-cells and sera from variant BAC-Tg mice phenocopy cytokine excess observed in APOL1-RV patient sera following viral stimulation.** (A-F) Splenic percentages of (A) CD4+, (B) CD8+ T-cells, (C) B cells, (D) NK cells, (E) monocyte-macrophages, and (F) dendritic cells (G) Tregs at day 7 following injection with poly(I:C) in G1- and G0-hybrid mice. (H) Heat map of differential serum cytokines expression from day 0 (day of PCR positivity of SARS-CoV-2 infection) to day 28 observed in the CHARM cohort with respect to their demographics (age, sex, and race), APOL1 genotype (either G1/G2, G1/G0, G2/G0, or G0/G0), and the symptoms (none, mild and moderate). (I-K) Line graphs of serum levels of (I) TNFa, (J) IL-2, and (K) CXCL6 in G1 or G2-carrying patients vs G0 before from day of PCR positivity for SARS-CoV-2 infection to day 28. (L) Each bar represents significantly elevated serum cytokine levels observed in COVID-19 sera in the CHARM cohort overlapping with cytokines also upregulated in CD8+T-cells from variant APOL1-BAC-Tg mice. The bar graphs (A-G) display mean ± SEM. *$p < 0.05$; circles/triangles represent individual mice.

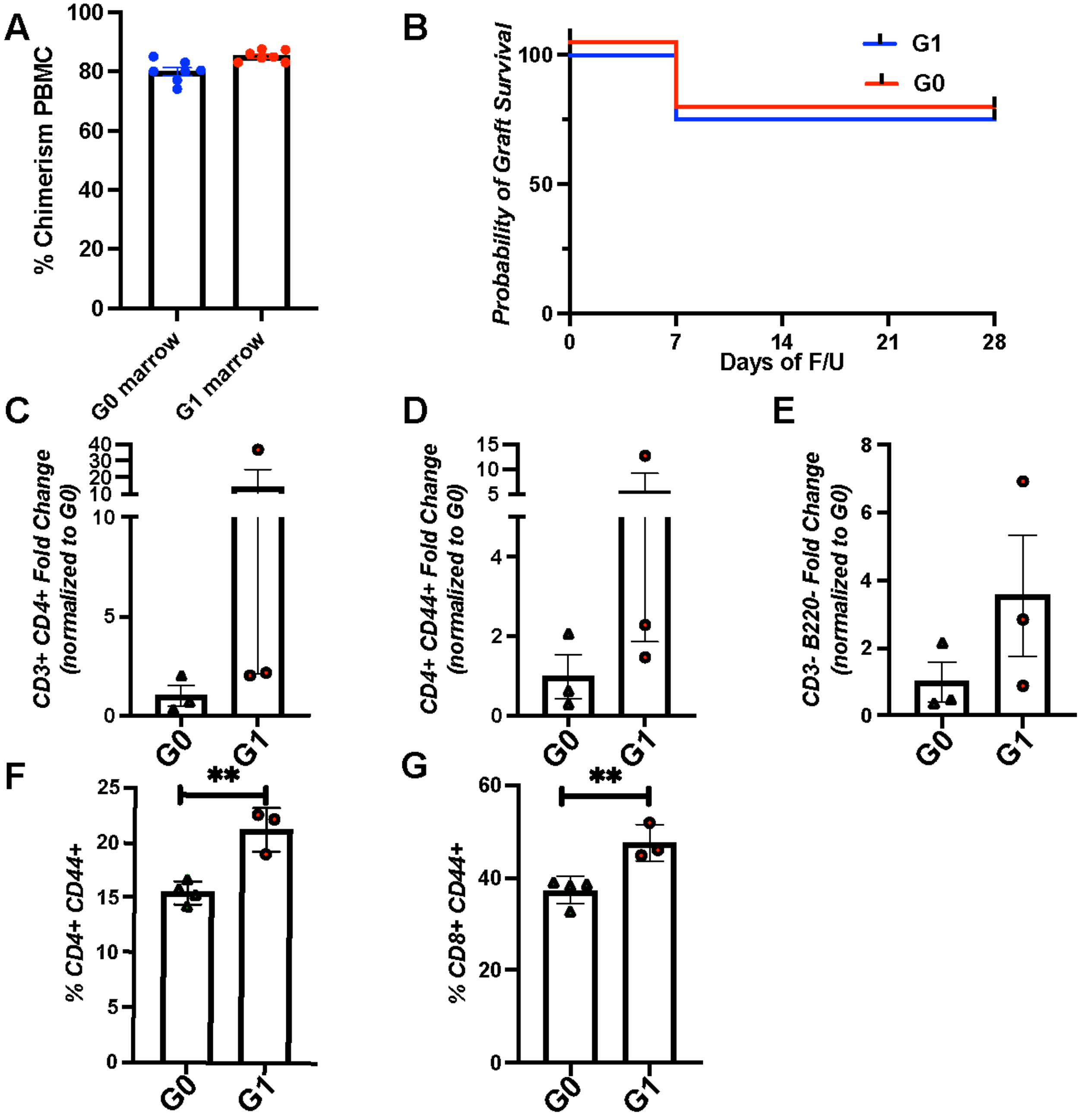

**Supplemental Figure 6: Variant APOL1-BAC-Tg mice demonstrate accelerated allograft rejection in an allogenic heart transplant model.** (A) Chimerism of Bl6/45.2-G0- and G1-bone marrow mice were confirmed by measuring CD45+45.2+ cells in PBMC at 4 weeks in chimeric Bl6/45.1 hosts. (B) Kaplan-Meier curve of allograft survival data in G1-chimeric recipients vs G0 controls without sensitization; 3/4 of allografts survived over 28 days. (C-E) FACS analysis of allograft infiltrating cells at day-28 shows the proportion of (C) CD4+, (D) CD4+CD44+, and (E) CD3-B220- (innate immune) cells in G1-chimeric recipients vs G0. (F-G) FACS analysis of splenocytes obtained at day-7 from a subset of sensitized G1- and G0-chimeric recipients and allotransplanted mice with proportions of (F) CD4+CD44+ and (G) CD8+CD44+T-cells in G1-chimeras vs G0-chimeras. The bar graphs display mean ± SEM. $^{**}p < 0.01$, circles/triangles represent individual mice.

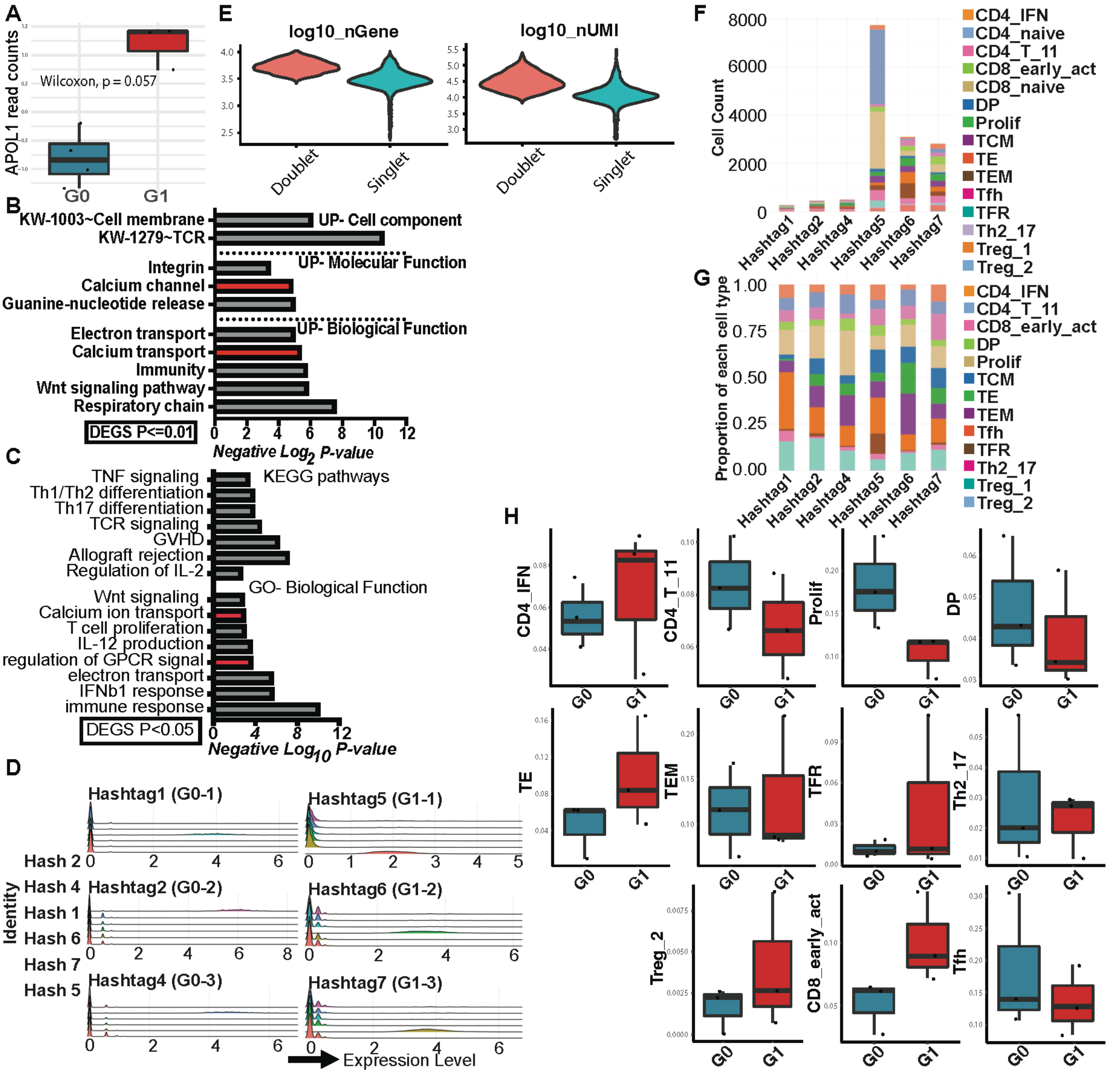

**Supplemental Figure 7: Transcriptome analyses from BAC-Tg CD8+ T-cells relays a role for enhanced Calcium-mediated Calcineurin/NFAT signaling in G1-CD8+T-cells.** (A) Expression of *APOL1* in stimulated CD8+T-cells (expressed as transcript per million normalized by library size) in each genotype. (2-sided Wilcoxon Rank Sum Test) (B-C) Functional enrichment of DEGs obtained (G1 vs G0) with (B) P < 0.01, (C) and P < 0.05 using DAVID (data base for annotation, visualization and integrated discovery). (D) Expression of the hashtag oligos in each sample of graft-infiltrating CD3+ cells (log-ratio (CLR) transformation). (E) The distribution of log10 (gene number) and log10 (UMI number) of cells identified as doublets and singlets. (F) The numbers of infiltrating T-cells of each subset identified from the six samples. G1-Sample#5 is an outlier with high numbers of naïve T-cells, G1-recipient grafts had overall greater T-cell infiltration. (G) The proportion of each cell type (after excluding naïve T-cells) within each sample is shown. (H) Box plots compare infiltrating proportions of selected T-cell subsets between 2 genotypes (G-H: Naïve cells excluded from proportion analysis due to outlier Sample#5).

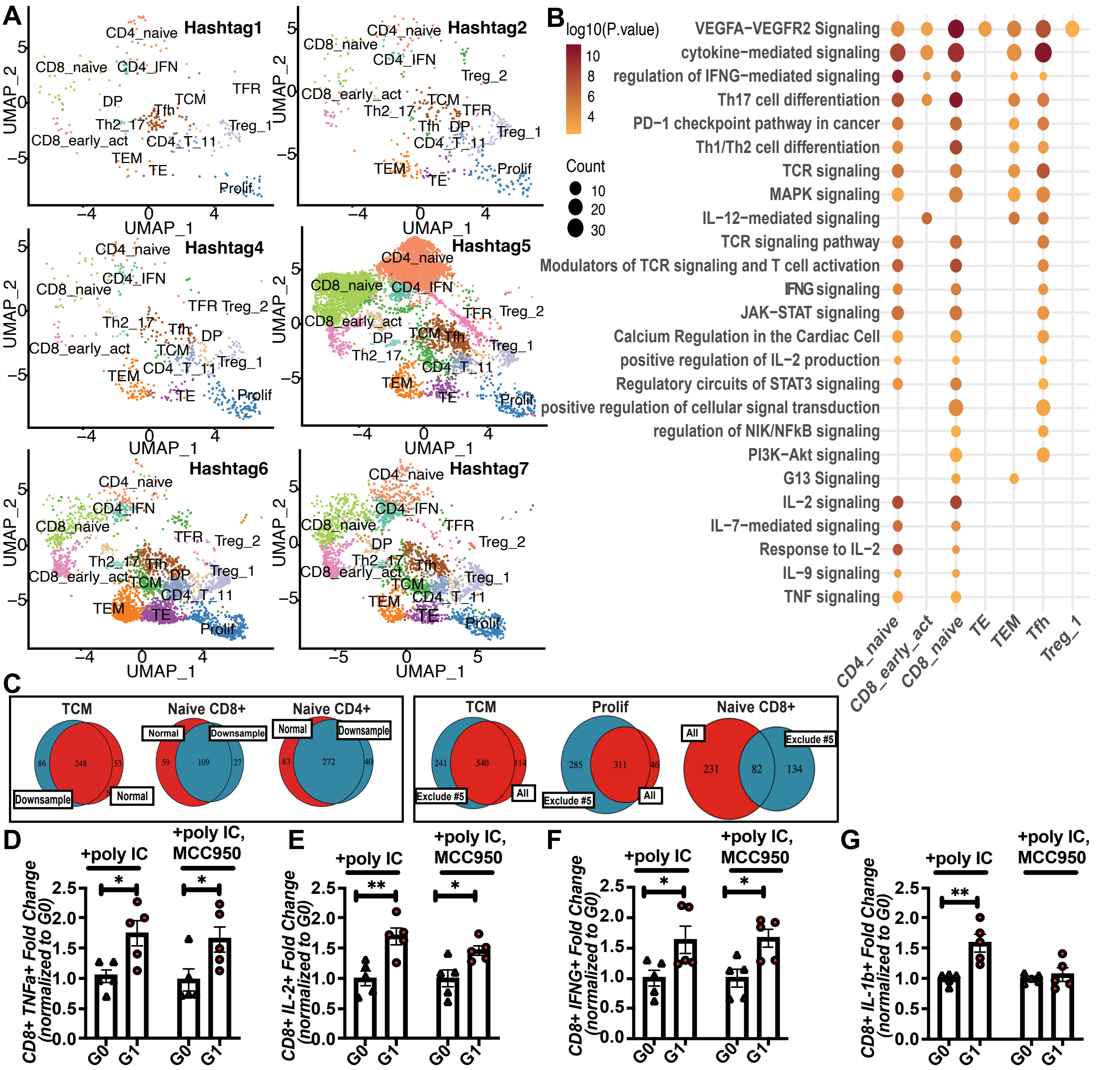

**Supplemental Figure 8: Transcriptome analyses from BAC-Tg CD8+ T-cells relays a role for enhanced Calcium-mediated Calcineurin/NFAT signaling in G1-CD8+T-cells (contd.)** (A) UMAP of cell populations in each of the 6 samples (n=3 each per genotype). (B) Bubble plots show enriched functions of the DEGs (G1 vs G0; P<0.01) obtained from each cell type is shown. (C) Venn diagrams show overlap of DEGs from all 6 samples, vs those identified after excluding G1-Sample#5 [left panel], and similar findings comparing downsampled DEGs vs directly obtained (including all cells; see Methods) [right panel] for TCMs, CD8+ naïve T cells and CD4+ naïve T-cells (hypergeometric P<0.001 for each). (D-G) CD8+T-cell-specific relative production (expressed as fold change and normalized to G0) of (D) TNFa, (E) IL2, (F) IFNG, and (G) IL-1b in ex vivo poly (I:C)-treated splenocytes with or without MCC950. The bar graphs display mean ± SEM. *p < 0.05, **p < 0.01.

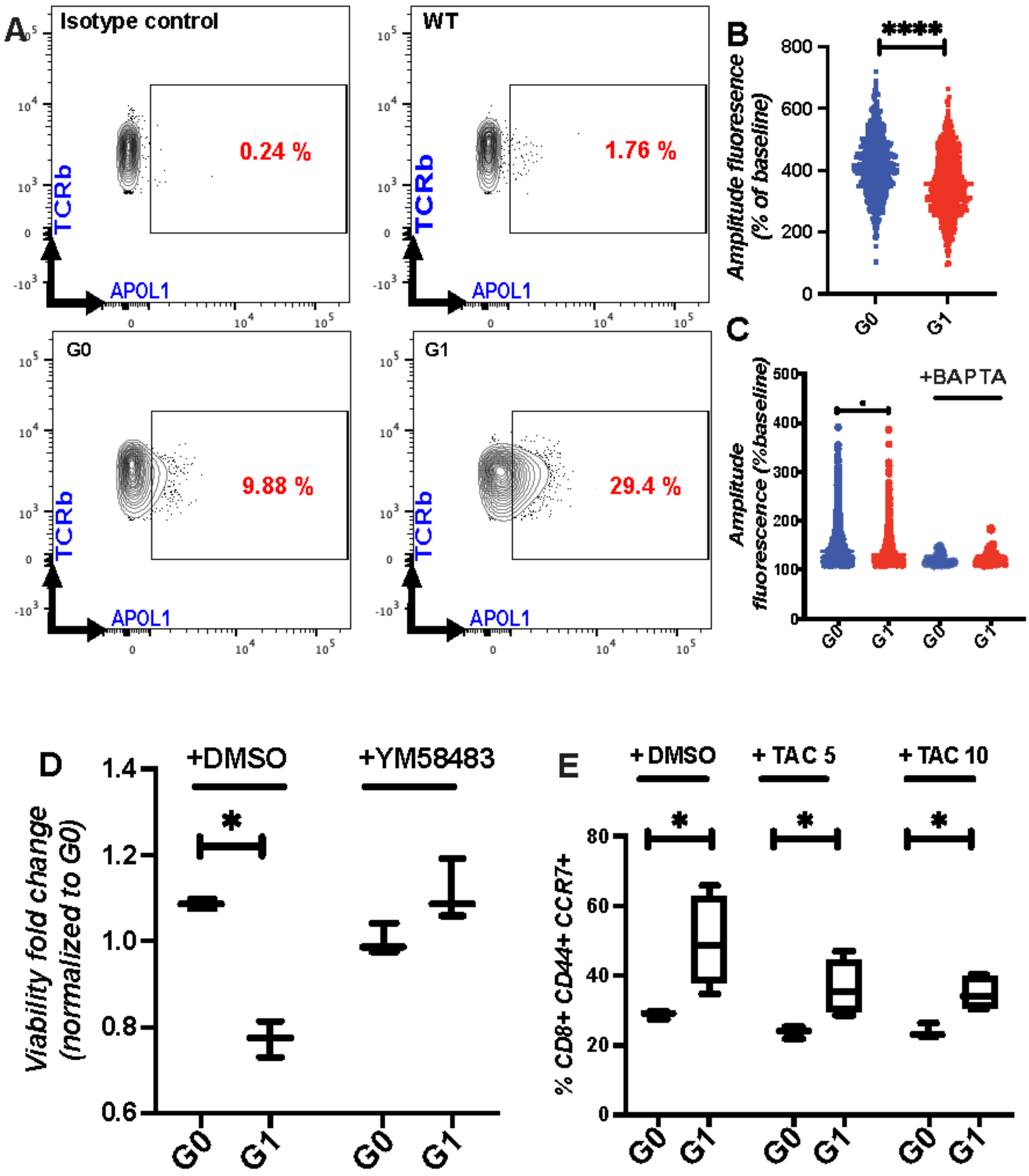

**Supplemental Figure 9: ER calcium depletion causes activation of variant APOL1 CD8+T-cells.** (A) Representative flow plots of APOL1 in G1 with an isotype control, WT, G0, and G1, showing surface expression. (B-C) CD8+T-cells were isolated and stimulated in 96-well plates with anti-CD3/CD28, and live cell imaging was performed. (B) Amplitudes of -fluorescence measured from live G0- and G1-CD8+T-cells in response to Thapsigargin [N=5 mice each], while (C) shows mean amplitude of fluorescence from CD8+T-cells from G1 vs G0 after anti-CD3-stimulation. BAPTA-AM completely abolished fluorescence [(>500 cells, n>6 mice]. (D) Viability of G1 CD8+ T-cells vs. G0 (fold-change) treated with YM-58483 at 50 nM and DMSO. (E) Dose-dependent effects of TAC treatment on TCM expansion in G1- and G0- naïve CD8+T-cells. Box-Whisker plots show median and range of triplicate wells pooled from 5 mice each. *p < 0.05, ****p<0.0001. TAC: Tacrolimus.

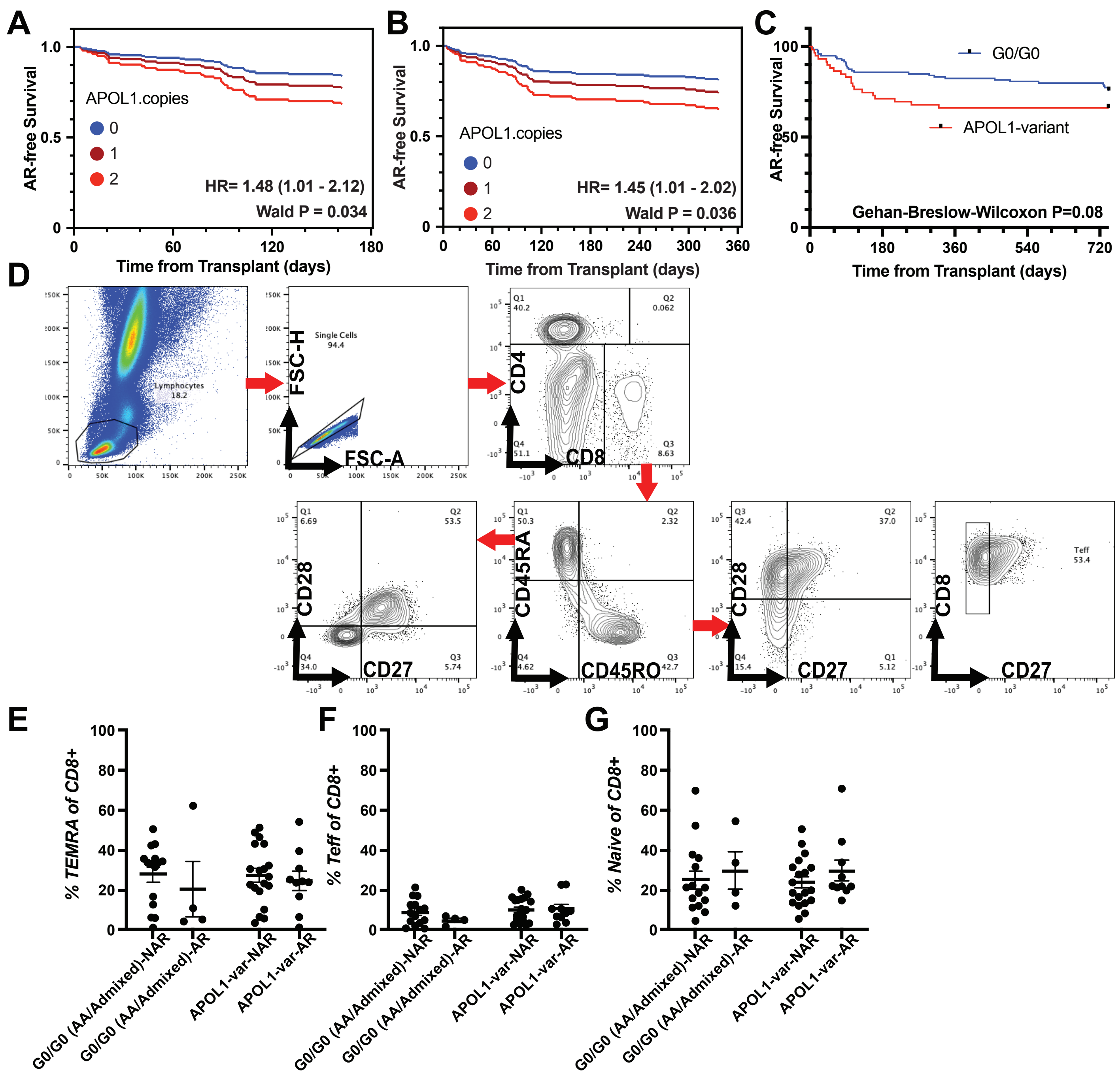

**Supplemental Figure 10: Association of graft rejection with APOL1 copy number and CD8+T cell phenotypes in CTOT-19 cohort.** (A-B) Time-to-event analyses of AR with APOL1 copy numbers in an additive model are plotted. The additive models show a positive association between the copy number addition and rejection (A) by 6 months (hazard ratio=1.48, Wald p=0.034) and (B) by 12 months (hazard ratio=1.45, Wald p=0.036) time point. (C) Time-to-event analyses of the rejection rate of APOL1 risk variant-expressing recipients vs. G0/G0 by 24 months (Wilcoxon p=0.08) (D) The gating strategy applied to PBMCs gated for TCM (CD8+CD45RO+CD28+CD27+), TEMRA (CD8+CD45RA+CD28-CD27-), T effector CD8+CD45RO+CD27-), and naive CD8+Tcells (CD8+CD45RA+CD28+CD27+). Pre-transplant proportions of (E)TEMRA (F) T effector and (G) naïve CD8+T-cells in patients with AR or NAR categorized by APOL1 genotypes are plotted.

Supplementary Table 6: Patients with biopsies used for AR/NAR analyses by APOL1-RVs in CTOT-19

| **Biopsy, N (%)** | **APOL1 G0/G0, N (%)** | **APOL1-RVs, N (%)** | **P Value (<0.05)** |
|---|---|---|---|
| 6 Months Central (yes) | 96 (66.2) | 36 (55.3) | 0.274 |
| 6 Months Local (yes) | 114 (78.6) | 55 (84.6) | 0.351 |
| 24 Months Central (yes) | 102 (70.3) | 40 (61.5) | 0.264 |
| 24 Months Local (yes) | 118 (81.3) | 59 (90.7) | 0.101 |

***Note:** Supplemental tables 1-5 in attached Supplemental Data Excel

Supplementary Table 7: Adjusted Cox proportional hazards models testing APOL1-variant genotype and AR

| **Model-1** | **Hazard ratio** | **95% CI** | **P Value** |
|---|---|---|---|
| Recipient age | 0.87 | 0.9554 to 1.018 | 0.38 |
| Any.APOL1 Variant [Ref G0/G0] | 2.083 | 1.042 to 4.161 | **0.03** |
| **Model-2** | | | |
| CMV. Intermediate risk [Ref D-/R-] | 0.9912 | 0.3540 to 3.520 | 0.98 |
| CMV. Intermediate risk [Ref D-/R-] | 1.045 | 0.3065 to 4.074 | 0.94 |
| Any.APOL1 Variant [Ref G0/G0] | 2.219 | 1.091 to 4.565 | **0.02** |
| **Model-3** | | | |
| Infliximab Limb [Ref: Placebo] | 0.6593 | 0.3298 to 1.295 | 0.22 |
| Any.APOL1 Variant [Ref G0/G0] | 2.222 | 1.127 to 4.381 | **0.02** |
| **Model-4** | | | |
| AA Ancestry [Ref: European] | 0.9138 | 0.2591 to 2.888 | 0.88 |
| Other Ancestry [Ref: European] | 1.063 | 0.3467 to 2.956 | 0.90 |
| Any.APOL1 Variant [Ref G0/G0] | 2.391 | 0.8683 to 7.386 | 0.10 |

Supplemental Table 8: List of primers

| | | |
|---|---|---|
| Human | APOL1 | Copy number |
| | F | CCGGGTCACTGAGCCAATC |
| | R | ACACGAGGTAGACTACATCCAG |
| Human | APOL1 | mRNA |
| | F | CCGGGTCACTGAGCCAATC |
| | R | ACACGAGGTAGACTACATCCAG |
